# Divergent lineage trajectories and genetic landscapes in human gastric intestinal metaplasia organoids associated with early neoplastic progression

**DOI:** 10.1101/2024.04.03.588024

**Authors:** Sarah SK Yue, Yin Tong, Hoi Cheong Siu, Siu Lun Ho, Simon YK Law, Wai Yin Tsui, Dessy Chan, Yuanhua Huang, Annie SY Chan, Shui Wa Yun, Ho Sang Hui, April S Chan, Siu Tsan Yuen, Hans Clevers, Suet Yi Leung, Helen HN Yan

## Abstract

**Objective:** Gastric intestinal metaplasia (IM) is a pre-cancerous stage spanning a morphological spectrum that is poorly represented by human cell line models. We aim to establish and characterize human IM cell models to better understand IM progression along the cancer spectrum.

**Design:** A large human gastric IM organoid (IMO) cohort (n=28) was established, along with normal gastric organoids (n=42) for comparison, and comprehensive multi-omics profiling and functional characterization were performed.

**Results:** Single-cell transcriptomes revealed IMO cells spanning a spectrum from hybrid gastric/intestinal to advanced intestinal differentiation, and unveiled lineage trajectories that connected different cycling and quiescent stem and progenitors, highlighting their differences in gastric to IM transition. The hybrid IMO cells showed impaired differentiation potential, high lineage plasticity beyond gastric or intestinal fates, and reactivation of a fetal gene program.

Cell populations in gastric IM and cancer tissues were found to be highly similar to those derived from IMOs and exhibited fetal signature. Genomically, IMOs showed an elevated mutation burden, frequent chromosome 20 gain, and epigenetic de-regulation of many intestinal and gastric genes. Functionally, IMOs downregulated *FGFR2* and became independent of FGF10 for survival. Several IMOs exhibited a cell-matrix adhesion independent (CMi) subpopulation that displayed chromosome 20 gain but lacked key cancer driver mutations, which could represent the earliest neoplastic precursor of IM-induced gastric cancer.

**Conclusions:** Overall, our IM organoid biobank captured the heterogeneous nature of IM, revealing mechanistic insights on IM pathogenesis and its neoplastic progression, offering an ideal platform for studying early gastric neoplastic transformation and chemoprevention.

## Significance of this study

### What is already known on this topic?

Gastric IM is histologically classified into incomplete and complete forms, and confers an increased risk of gastric cancer (GC).

Molecular profiling of IM tissues has revealed an elevated somatic mutation rate, new IM drivers, chromosomal alteration, telomere shortening and aberrant DNA hypermethylation.

Single-cell RNA sequencing of IM tissues showed a gastric-intestinal mosaic phenotype, revealing the presence of IM stem cells and uncovering some novel pro-tumorigenic fibroblast subtypes.

Given the complexity and existence of different stem/progenitor populations in gut homeostasis and injury repair, detailed information on IM stem/progenitor populations remains limited.

Although some studies have used IM organoid models for cell biology research, there is a lack of models with detailed phenotypic characterization across a wide histological spectrum.

### What this study adds?

We have established a large series of IM organoids, covering the heterogeneity of IM and spanning various grades of gastric to intestinal transition.

Single-cell transcriptomic analysis of our stem/progenitor enriched gastric and IM organoids revealed the lineage trajectories of several IM-specific stem/progenitors that were previously not well-characterised, and displayed high concordance with cellular subgroups found in IM and GC tissues.

We discovered that hybrid IM cells, being trapped in mixed gastric and intestinal differentiation, displayed lineage plasticity and fetal reversion. Such a phenotype could facilitate cancer progression and indeed was observed in gastric cancer tissue.

We found gain of FGF10 independence in the metaplastic process through downregulation of FGFR2 in IMOs.

Our IMO models captured a wide spectrum of genetic and epigenetic perturbations, highlighting a unique model of IM progression through chromosome 20 gain, with phenotypic diversity adaptation to achieve anchorage independent growth.

### How this study might affect research, practice or policy?

The gastric and IM organoid dataset can be further explored together with tissue datasets to comprehensively decipher stem cell biology in IM and cancer initiation.

The IM organoid models complement existing cancer models and can be used for investigating early gastric neoplastic transformation and prevention.

The advanced cell-matrix adhesion independent IM cells with chromosome 20 gain might mark early malignant cells, potentially aiding the identification of high-risk IM patients with potential for gastric cancer development.

## INTRODUCTION

Gastric intestinal metaplasia (IM) is a pre-cancerous stage of gastric cancer (GC) characterized by the transformation of gastric cells into a more intestinal-like cell type. This process is driven by the ectopic expression of transcription factors *Cdx1* or *Cdx2* in the gastric mucosa of transgenic mice, leading to the development of IM.^1–3^

The prevalence of IM and GC varies globally, with East Asia, particularly China, exhibiting high incidence rates.^4^ ^5^ Various methods have been developed to classifying IM in an effort to predict the risk of GC. Traditional histochemical methods that categorized IM into complete and incomplete types were previously suggested to have different implications for GC risk, but these classifications are no longer widely accepted.^5^ Subsequent molecular studies have revealed the presence of p53 mutations,^6^ ^7^ microsatellite instability^8^ and epigenetic deregulation^9^ in IM tissues associated with GC progression. More recently, multi-omics studies in IM tissues have identified increased somatic mutation rates, new IM drivers, chromosomal alterations, telomere shortening and aberrant DNA hypermethylation.^10–13^ Long-term follow-up studies have suggested that data on chromosomal alterations, telomer shortening, and the integration of transcriptomic and genomic information with clinical data could stratify IM and identify patients at higher risk of GC progression.^10^ ^11^ As a subset of GCs have been shown to share a transcriptional signature or epigenomic footprint with IM,^13–15^ detailed characterization of IM cells may provide insights into their potential for neoplastic progression. Indeed, single-cell RNA sequencing (scRNAseq) of IM tissues has identified IM cells with a gastric-intestinal mosaic phenotype and discovered some novel pro-tumorigenic fibroblast subtypes.^11^ ^16–19^ Although epithelial stem cells are generally under-represented in scRNAseq studies due to their inherent low numbers in tissue and also because they are affected by the prolonged tissue digestion process, recent studies have shown the presence of putative IM stem cells with expression of markers such as *OLFM4* or *LEFTY1*.^11^ ^18^ ^19^ However, detailed information on various subpopulations of stem and progenitors in the gastric to IM transition remains limited. Given the heterogeneity of the IM process, a viable cell model for IM that covers a wide spectrum of phenotypes would be invaluable for the development of novel strategies for GC prevention. It can also provide information on IM stemness, lineage plasticity, differentiation trajectories and signaling pathway deregulation, thereby potentially shedding additional light on why IM confers susceptibility and propensity for neoplastic progression.

In this study, we established a large gastric IM organoid cohort derived from GC patients. We conducted comprehensive molecular profiling of the genomic, epigenomic and transcriptomic alterations in these IM organoids, as well as performed scRNAseq to characterize stem/progenitor populations and their corresponding lineage trajectories. We discovered lineage plasticity in IM beyond gastric or intestinal fates, and demonstrated a reversion to a fetal phenotype. Through in-depth functional characterization of the IM organoids, we identified several subpopulations of IM cells with cell-matrix adhesion independent (CMi) capacity. We further identified a link between chromosome aberration, the intestinal differentiation program and CMi capacity in IM organoids, potentially working in concert to convey one of the earliest tumour initiating events for GC development.

## METHODS

See online supplemental methods for details on the study patient cohort (online supplemental figure 1), organoid culture conditions, various cell based assays, immunoblotting, immunohistochemistry, DNA/RNA extraction, qRT-PCR, droplet digital PCR, whole-exome sequencing, RNA sequencing, scRNAseq, single-cell CNV (scCNV) profiling, methylation profiling and various bioinformatic analyses.

## RESULTS

### Establishment of an IM organoid cohort and evaluation of the transcriptomic landscape

Previously, we established a gastric and colon cancer organoid biobank, which included linked genomic, transcriptomic and patient clinico-pathological data.^20^ ^21^ Here, we expanded this biobank by establishing a collection of 70 organoids derived from normal gastric mucosa and gastric tissues with IM from 47 GC patients. Initially, we classified normal-IM paired adjacent frozen tissue according to the extent of IM, as determined histologically by a pathologist, and the organoids derived from them were given the designation “IMO”, indicating organoids with moderate to extensive IM, or “BO” or “AO”, indicating normal organoids derived from gastric body or antrum tissue, respectively, with no or only mild (focal) IM. The clinico-pathological parameters for each sample are summarized in online supplemental table 1 and sequencing statistics for those that were included for multi-omics analysis are summarized in online supplemental table 2.

Unsupervised hierarchical clustering of RNAseq data from the 70 organoids revealed two distinct clusters (figure 1A). The majority of AOs and BOs clustered together, while most of the IMOs clustered with colon/ileum organoids (COs), albeit in two separate groups (IMOexp cluster 1 and IMOexp cluster 2). Although both IMO clusters showed upregulation of intestinal lineage markers (green bracket), the downregulation of the gastric lineage markers (red bracket) was more pronounced in cluster 1. This could be due to a high purity of IM cells in cluster 1, along with an advanced shift in the transcriptome towards more intestinal-like cells, or both (as detailed in single-cell transcriptome analysis below). Based on the transcriptome data, we reclassified 8 organoid samples, including 4 organoids derived from mild IM tissues (GX035-AO, GX091-AgO, GX091-BlO and GX092-BO) that we originally classified as normal based on histology but clustered with IMOs, and 4 organoids derived from moderate IM tissues (GX080-IMO, GX086-IM2O, GX087-IM2O and GX093-IM2O) that were originally classified as IM but clustered with AOs/BOs. This suggests that a subpopulation of IM or gastric cells became dominant during the organoid culture process, thus requiring the samples to be re-classified. The final re-classified cohort consisted of 28 IMOs, 23 BOs and 19 AOs.

**Figure 1.**
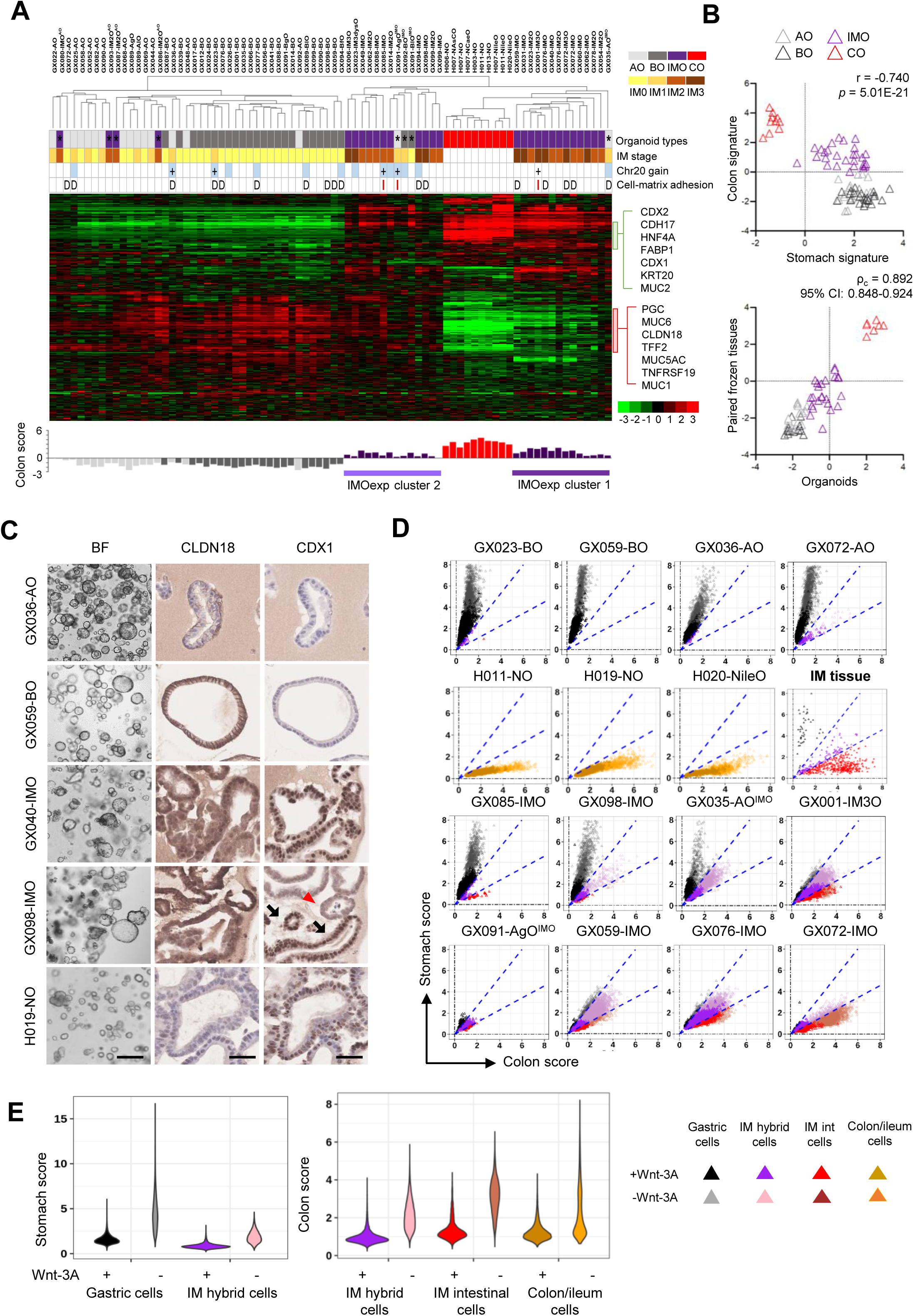
Generation of an IMO cohort with distinct transcriptomic profiles and bi-lineage cellular composition. (A) Hierarchical clustering using bulk RNAseq data. Heatmap (top) shows differential gene expression based on the mean-centered log2 TPM values of each gene (top). Organoid types correspond to the tissue types from which they were established. ‘*’ indicates reclassified organoids. IM stage denotes the extent of IM, as graded from the paired frozen tissues. Organoids with chromosome 20 gain, inferred by WES or by scRNAseq data, are denoted in blue or indicated by ‘+’, respectively. Regarding the cell-matrix (CM) adhesion dependence of the organoids, ‘I’ denotes independent, ‘D’ denotes dependent. Plot (bottom) showing positive colon signature scores for IMOs and COs, and negative scores for gastric organoids. (B) Scatter plot (top) showing a negative correlation between the human stomach and colon signature scores for the 4 different organoid types. Scatter plot (bottom) showing high concordant lineage specificity between organoids and their paired frozen tissues in terms of the aggregate score. (C) Representative brightfield (BF) and IHC images of the organoids. IMOs showed heterogeneous CDX1 expression, with positive staining indicated by black arrows and negative staining indicated by red arrowheads. Scale bars: 500µm (BF) and 50µm (IHC). (D) Scatter plots showing the human stomach and colon signature scores for each cell by scRNAseq in sample pairs of organoids with and without Wnt-3A. Each dot represents one cell and is coloured according to cell type classification based on the stomach and colon signature scores (see colour legend in (E). (E) Violin plots showing the extent of increase in both stomach and colon signature scores as a reflection of their differentiation upon Wnt withdrawal among different cell subgroups. The extent of differentiation in IM-hybrid cell was lowest compared to gastric and colon/ileum cells, and also lower than IM-Int cells denoting IM with advanced intestinal differentiation. (A-E) BO/BfO/BgO/BlO, body organoids; AO/AgO, antrum organoids; CO, intestinal organoids; IMO/IM2O/IM3O/IM3dysO, IM organoids.

To verify our organoid classification, we used stomach and colon gene signatures to score each organoid sample (online supplemental table 3). We observed a negative correlation between stomach and colon signatures for the different organoid types, with COs having high colon and low stomach signature scores, AOs/BOs having low colon and high stomach signature scores, and IMOs having intermediate scores for both signatures (r = -0.740, *p* = 5.01E-21) (figure 1B). An aggregate score was further derived for each sample, which showed a high concordance correlation between the frozen tissues and organoids (ρ_c_ = 0.892, 95% CI: 0.848-0.924) (figure 1B). Moreover, the gene expression patterns between individual IMO-frozen tissue pairs were highly correlated (online supplemental figure 2), demonstrating that the organoids can recapitulate the gene expression of *in vivo* tissues.

Strong protein expression of CLDN18 was observed in the cell membrane of normal gastric organoids, while CDX1 was only weakly or not expressed (figure 1C and online supplemental figure 3). In contrast, intestinal organoids exhibited strong expression of CDX1 in cell nuclei and absence of CLDN18 expression. Most IMOs appeared cystic and morphologically indistinguishable from normal gastric and intestinal organoids. However, they were characterized by strong nuclear expression of CDX1, and varying degrees of weakened membranous CLDN18 expression. Although a large population of CDX1^+^ cells were detected in organoid samples with high IM content (e.g. GX040-IMO), others (e.g. GX098-IMO) displayed a heterogeneous CDX1 staining pattern, which suggests a mixed population of IM and normal gastric cells.

### Single-cell transcriptomic analysis reveals the distinct cellular composition and bi-lineage differentiation potential of IMOs

To dissect the cellular composition and lineage plasticity of IM compared with gastric cells, we cultured the organoids with or without Wnt-3A to encourage stem cell enrichment or induce differentiation, respectively, followed by scRNAseq. Their associated scRNAseq statistics encompassing 92,730 cells are summarized in online supplemental table 4.

The cellular composition of each organoid pair, cultured with or without Wnt-3A, was measured relative to the human stomach and colon signature scores of each single cell (figure 1D and online supplemental table 5). As expected the gastric organoids predominantly consisted of gastric cells (>97.5%), displaying a high stomach and very low colon signature score, while the intestinal organoids, derived from normal colon or ileum, were primarily composed of intestinal cells (>97.4%), displaying a low stomach and high colon signature score. Upon differentiation (-Wnt-3A), they displayed even higher organ specific scores (figure 1D, E). Interestingly, the IMOs mainly consisted of hybrid cells expressing both the gastric and colon signatures (hereby named IM-hybrid), with a subpopulation showing more advanced intestinal differentiation and a marked reduction of in gastric scores, mimicking the intestinal cells (hereby named IM-Int) (figure 1D, E). For example, GX001-IM3O, GX091-AgO^IMO^, GX059-IMO, and GX076-IMO primarily contained IM-hybrid cells (88.1% to 93.3%), while GX072-IMO mostly contained IM-intestinal cells. Other IMOs (GX085-IMO, GX098-IMO, GX035-AO^IMO^) contained gastric cells in addition to IM-hybrid cells, likely due to an admixture of normal gastric glands in the original sample. Notably, these three cell lineage types were also present in fresh IM tissue (9.52% gastric, 21.6% IM-hybrid and 69.9% IM-int cells), confirming that our organoid culture model effectively captured these different IM states. Indeed, this hybrid cell status of IM cells observed *in vitro* has been recently identified in both gastric and Barrett’s esophagus IM cells *in vivo*,^11^ ^16–19^ further reinforcing that IM organoids accurately recapitulate tissue characteristics. Contrary to intestinal or gastric cells, which exhibited a significant increase in colon or gastric signature scores, respectively, upon differentiation induction, IM-hybrid cells showed a notable reduction in differentiation propensity, as demonstrated by only a modest increase in both the stomach and colon signature scores (figure 1E). A possible explanation could be that IMOs arrested in a hybrid state could have impaired differentiation ability.

Next, we constructed a gastric cell atlas by integrating gastric and IM organoid samples, identifying 36 unique epithelial cell clusters (online supplemental methods, online supplemental figure 4A). Based on the expression of marker genes that characterize specific gastric or intestinal cell populations, these clusters were merged into 14 cell groups representing various stem, progenitor and differentiating cell populations from gastric or IM samples (figure 2, online supplemental figure 4 B and C, online supplemental table 6). The relative distribution of these cell populations across samples, when cultured with or without Wnt3A, is shown in figure 2B. The expression of key stem cell and lineage specific markers in the various stem and progenitor populations of gastric and IMOs are shown in figure 2C. The differential gene expression of each cell population is provided in online supplemental dataset 1.

**Figure 2.**
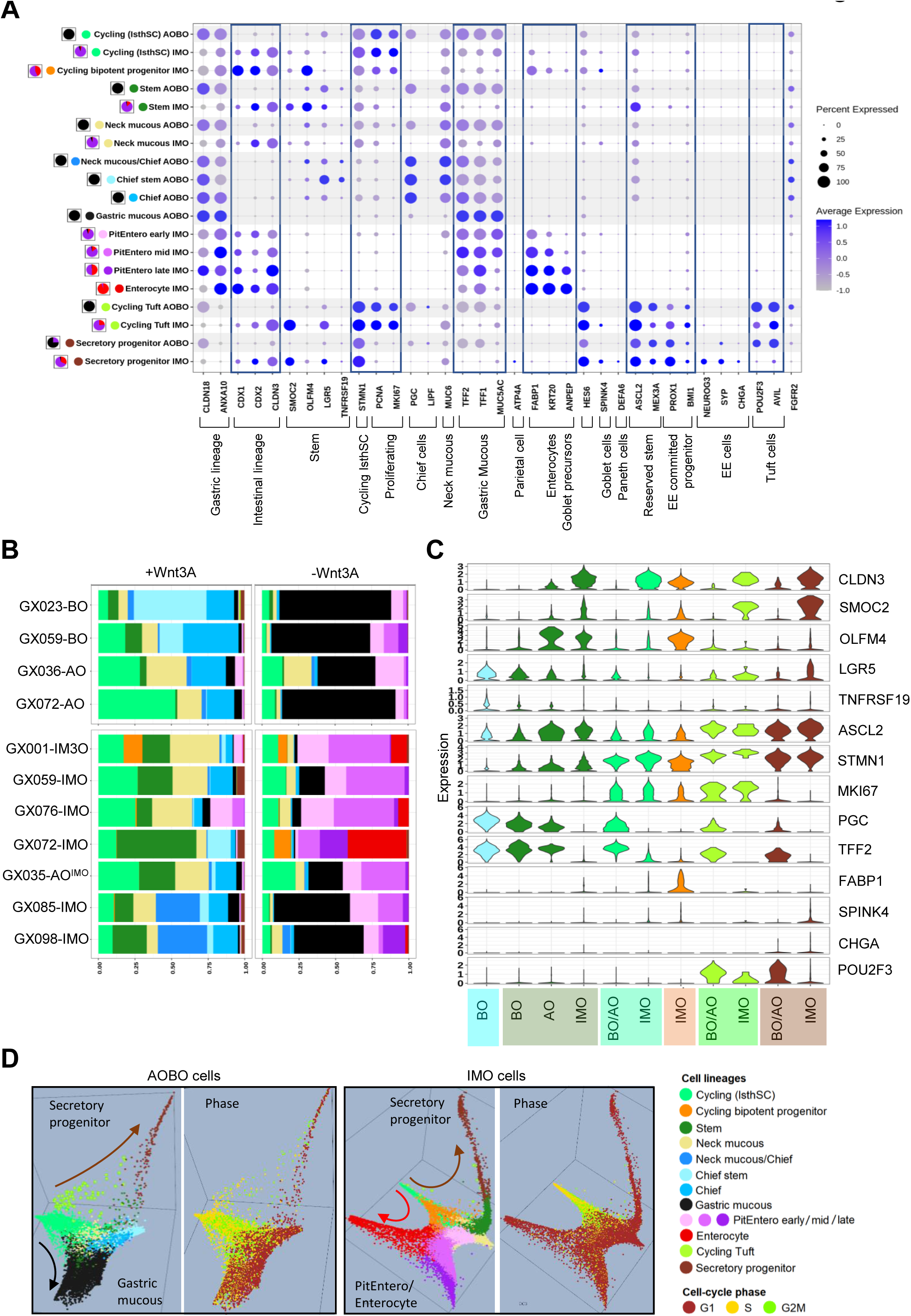
Single cell atlas of gastric and intestinal metaplasia organoids. (A) Dot plot of scaled expression of key marker genes grouped by cell type in 4 AO/BO gastric organoids (GX023-BO, GX059-BO, GX036-AO, GX072-AO) and 5 IMO samples with high IM purity (GX001-IM3O, GX091-AgO^IMO^, GX059-IMO, GX076-IMO, GX072-IMO), combining cells cultured with or without Wnt3A. Pie-charts on the left of each cell type denoted the fraction of gastric, IM-hybrid or IM-Int cells according to classification in figure 1D. The cell typing from all samples merged together can be found in online supplemental figure 4A-B. The colour denotes the mean expression level across the cells (dark blue indicates high expression), while the dot size indicates the proportion of cells that express the gene. AOs/BOs versus IMO are shown separately for each cell type. AO/BO data are shaded in grey in the background to distinguish from the IMOs. Result are shown for a group only if a specific cell type predominates in either AO/BO or IMO. (B) Stacked bar charts showing the cell type composition of each sample when cultured with (left) or without Wnt3A (right). (C) Violin plots comparing the expression of cell lineage markers in various populations of stem/progenitor cells by organoid type. The colours in the plots (B and C) indicate the cell type, as displayed in colour legend in D. (D) Diffusion map analysis with 3D cell lineage trajectory plots of AOBO and IMO cells, respectively. The top 3 diffusion components were plotted as the x, y and z axis. Cells are coloured according to cell lineage and cell cycle phase. AOBO cells showed cycling isthmus stem cells and cycling tuft cells, with differentiation gastric mucous cells and secretory progenitors. IMO cells showed three populations of cycling stem cells, including cycling bipotent progenitors, cycling isthmus stem cells and cycling tuft cells. They displayed two major differentiating trajectories towards secretory progenitors and PitEntero/enterocytes (See online supplemental figure 4D for additional panel).

We then projected the cell lineage differentiation trajectories of gastric and IM organoids in a 3-dimensional diffusion map based on RNA velocity-based cell-to-cell vectors and diffusion similarities, using an unsupervised time sequence method, CellRank^22^ (figure 2D, online supplemental figure 4D, online supplemental methods). This timeline revealed the chronological sequence of cellular differentiation between the single cells in both AO/BO and IMO samples. Analysis of the 3D diffusion map revealed two distinct differentiation trajectories originating from cycling isthmus stem cells (IsthSCs) in AOs/BOs, towards gastric mucous cells or secretory progenitors (figure 2D, online supplemental figure 4D). Similarly, in IMOs, the cycling IsthSCs or bipotent progenitors differentiated along two distinct trajectories, developing into PitEntero/Enterocytes and secretory progenitors. For clarity, we further investigated these two distinct trajectories separately in detail (figure 3 and online supplemental figure 5).

**Figure 3.**
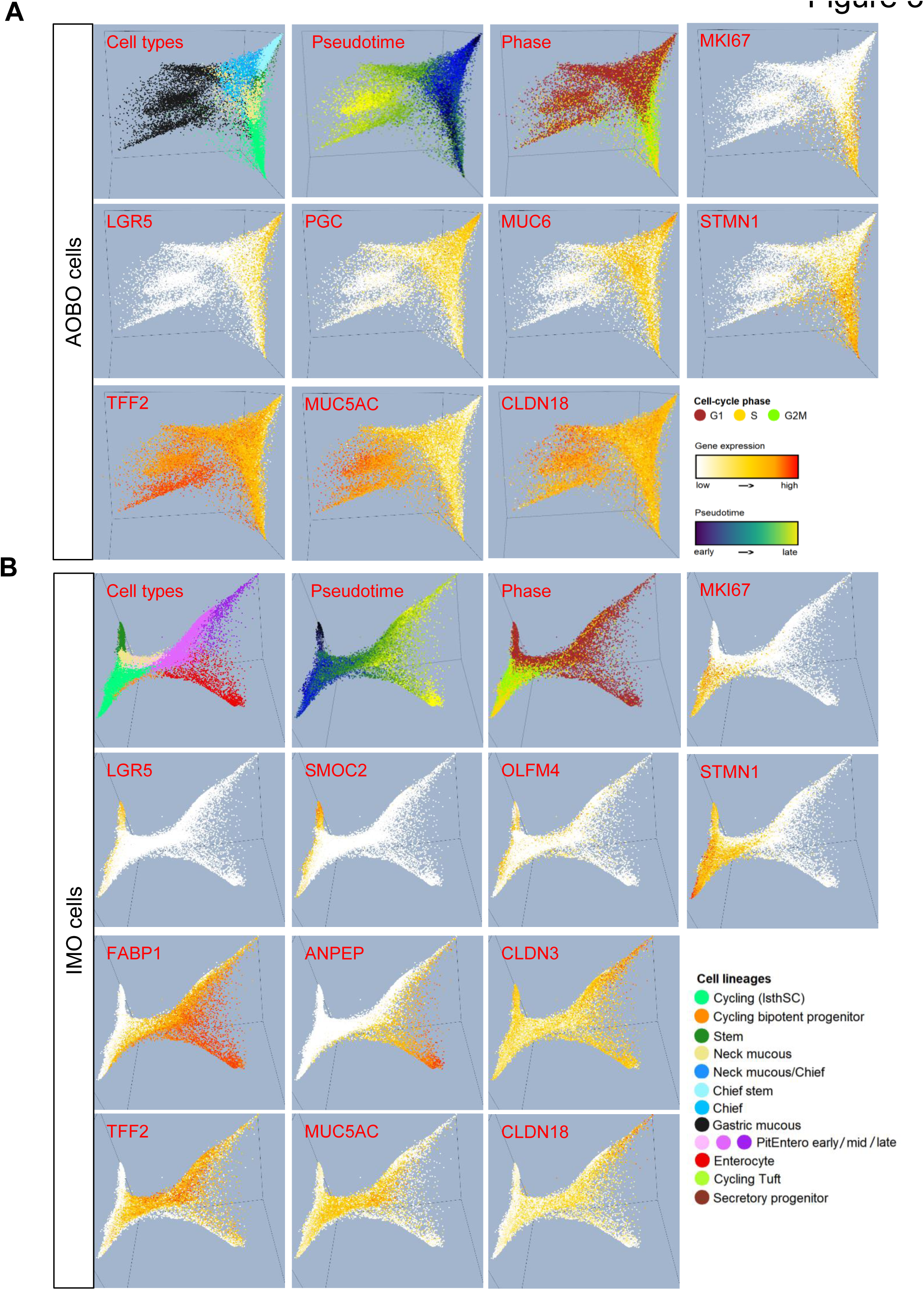
Gastric and IM cells showed distinct lineage trajectory. Diffusion map analysis with 3D cell lineage trajectory plots. Single cells from 4 AO/BO samples (with/without Wnt3A) were integrated into the AO/BO cell set (A), and single cells from 5 pure IMO samples (with/without Wnt3A) were integrated into the IMO cell set (B), after batch correction. The 3-dimensional diffusion map was plotted on the top 3 diffusion components as the x,y and z. Cells are coloured according to cell lineage, as denoted in right lower panel, pseudotime, cell cycle phase or by the expression intensity of a specific gene, as labeled in each box. Lineage trajectory towards gastric mucous for AO/BO and PitEntero/Enterocyte for IMOs are displayed here. IMOs showed expression of both gastric and intestinal genes, especially in the PitEntero cells. The IMO stem cells showed expression of intestinal stem marker *SMOC2*. See online supplemental figure 5 for the secretory lineage trajectory.

In mice, various subpopulations of gastric stem cells have been identified, including rapidly cycling isthmus stem cells (IsthSC) in the corpus, characterized by the expression of proliferation genes along with *stmn1*,^23^ actively cycling *lgr5* stem cells at the base of antral glands,^24^ and a quiescent stem cell population residing at the base of the corpus that express markers such as *lgr5* or *troy,* along with chief cell markers, acting as reserve stem cells.^25^ ^26^ However, less is known about human gastric stem cell populations. Here, we show that normal BOs/AOs showed cycling cell clusters with high expression of *STMN1, PCNA* and *MIK67,* but lower expression of other stem cell markers, closely resembling IsthSCs (figure 2A-C and figure 3A). Upon exiting the cell cycle, the cycling IsthSCs in the BOs/AOs transitioned towards various lineages of differentiation, including mucous neck cells (*MUC6* high), mucous neck/chief hybrid cells and chief cells (*PGC* high). Interestingly, we also observed a transition to a quiescent stem cell population specific to BOs, which we hereby refer to as chief stem cells, as they expressed *LGR5*, *TNFRSF19* and chief cell marker, *PGC*, resembling the corpus reserve stem cell population observed in mice (figure 2C). Upon Wnt-3A withdrawal, they differentiated into more mature gastric mucous cells (*TFF2*, *TFF1*, *MUC5AC* high). The BOs/AOs also showed a subpopulation of cells with secretory/enteroendocrine lineage differentiation trajectories, including a cycling tuft cell population (*POU2F3*, *AVIL*, *PCNA*, *MKI67* high) derived from the cycling IsthSCs, along with a quiescent tuft cell population (*POU2F3*, *AVIL* high; *PCNA*, *MKI67* low) derived from the quiescent *LGR5* stem cells. Both populations expressed the reserve stem cell markers *ASCL2* and *MEX3A*, along with EE progenitor markers *PROX1* and *BMI1* (Figure 2A and online supplemental figure 5).

IMOs displayed two populations of cycling stem cells, including the cycling IsthSC population expressing high *STMN1*, similar to gastric organoids and enriched in IM-hybrid state, and an IM-specific cycling population (*PCNA*, *MKI67* high) expressing enterocyte marker *FABP1*, goblet cell markers *HES6* and *SPINK4*, along with intestinal stem cell marker *OLFM4*, and enriched in IM-Int state (figure 2 A-C, figure 3B). Thus, these cells mimic the bipotential early progenitor cells in the transient amplifying zone of the intestine before diverging into enterocyte or secretory progenitors. IM organoids cultured in Wnt-3A also displayed a higher proportion of quiescent stem cells compared to normal gastric organoids, expressing intestine-specific stem marker *SMOC2*, along with other stem cell markers such as *LGR5* and *OLFM4*, but with low levels of proliferation markers (*PCNA*, *MKI67*) (figures 2A, B and figure 3B). This pattern mirrors previously reported quiescent *LGR5* stem cells in the human colon.^27^ Figure 2C clearly shows the various stem/progenitor populations in BOs/AOs versus IMOs. IM quiescent or cycling stem/progenitor populations can be distinguished from their gastric counterparts by downregulation of the gastric-specific gene *TFF2,* and upregulation of intestinal gene *CLDN3*, even though the expression of other intestinal genes was not prominent in these undifferentiated states. Interestingly, intestinal-specific stem cell marker, *SMOC2,* was expressed in all IMO stem/progenitors whereas *OLFM4* was expressed in both IMO and AO stem cells, as well as IMO cycling bipotent progenitors. The data clearly demonstrate that the process of IM altered the nature of the gastric stem cells.

Upon Wnt 3A withdrawal, IMOs differentiated into hybrid cells expressing both gastric mucous (*TFF2, TFF1, MUC5AC*) and enterocyte markers (*FABP1, KRT20, ANPEP*), which we subsequently named PitEntero cells (figure 3B). These cells exhibited a spectrum of hybrid differentiation, characterized by decreasing levels of *MUC5AC* and increasing levels of mature intestinal markers *KRT20* and *ANPEP* as IM advanced (figure 2A). Accordingly, we categorized the PitEntero cells into early, mid and late stage, with the most predominant population arrested in the mid-stage in most IMO cells. The subpopulation of IMO cells with advanced intestinal differentiation, as mentioned earlier, displayed higher expression of more advanced intestinal maturation marker *ANPEP* and lower level of gastric markers (e.g. *TFF2*, *MUC5AC*, *CLDN18*), and were therefore referred to as enterocytes. They predominantly displayed the IM-int state.

The secretory lineage differentiation trajectories of IMOs revealed that the cycling bipotent progenitor gave rise to the *SPINK4*+ goblet cell lineage, while the enteroendocrine (EE) progenitors derived from quiescent *LGR5*+ stem cells and displayed high levels of the intestinal-specific stem marker *SMOC2* (Figure 2A, online supplemental figure 5). Similar to BOs/AOs, both cycling and quiescent tuft cells (*POU2F3*, *AVIL* high), along with various EE cells (*NEUROG3*, *SYP*, *CHGA*, *SCG5* high), shared expression of reserve stem cell and EE precursor markers *ASCL2*, *MEX3A*, *PROX1* and *BMI1*. Overall, the various subpopulations of EE progenitors and tuft cells captured in our organoid cultures closely resemble the lineage trajectories previously described in the mouse intestine based on lineage tracing experiments, which also demonstrated that these EE cells possessed injury-inducible stem cell activity with putative origin from the reserved stem cell at the +4 position.^28^

### IM hybrid cells display lineage plasticity and fetal stemness

A recently discovered hallmark of intestinal tumorigenesis is the reversion to a fetal gene program and a more plastic cell fate.^29^ ^30^ This key feature highlights the shift towards an earlier developmental state, which allows for increased cellular plasticity and may contribute to the initiation and progression of cancerous transformations within the intestinal tissue. Concordantly, lineage plasticity has been shown to be exacerbated when niche factors are deficient, as demonstrated in metastatic colon cancer and their corresponding organoid culture.^31^ We therefore explored the lineage plasticity and expression of the fetal program in the IMOs.

We constructed a cell fate tracking system (CFT) (online supplemental methods) to unbiasedly quantify cell lineage diversity and cell plasticity states. We then applied a cell fate weighted matrix based on cell-specific expression genes of 81 cell types obtained from the Human Protein Atlas (HPA) tissue cell collection,^32^ from which we were able to calculate the cell fate differentiation potentials of every single cell transcriptome in our dataset into these 81 cell types (online supplemental table 7). We then defined the cell lineage diversity score as the Shannon-entropy of the 81 cell fate potentialities. We found that the majority of the AO/BO cells had a differentiation timeline that converged into a single destiny of gastric cells (figure 4A). For the IMOs, the IM-int cells displayed more intestinal and less gastric fates, but also a higher lineage diversity score compared with the AOs/BOs. Interestingly, the IM-hybrid cells maintained a much higher cell lineage diversity score even towards the end time stage, expressing not just gastric and intestinal cell fates, but also aberrant epithelial fates of cells that do not belong to the gastrointestinal tract, and even basal squamous or mesenchymal cell fates. Representative genes for basal/squamous cells (*NMU*), mesothelial cells (*CHAC1*), non-gastrointestinal (GI) epithelial cells such as lung alveolar cells type II (*PCSK9*), renal collecting duct cells (*HOXB6*), and prostatic glandular cells (*RAB3B*) are shown with their relative levels of expression along the differentiation timelines, highlighting their elevated levels especially in IM-hybrid cells, particularly those arrested in the PitEntero stage (figure 4C, D). We also used Waddington’s model^33^ to visualize cell-time-fate dimensions as a rugged landscape of sloping valleys and ridges, showing pluripotent cells rolling down into valleys of terminal differentiation (figure 4B). The IM cells, especially the IM-hybrid cells, displayed persistent high lineage plasticity in this landscape, with higher mountain altitude compared with the gastric cells.

**Figure 4.**
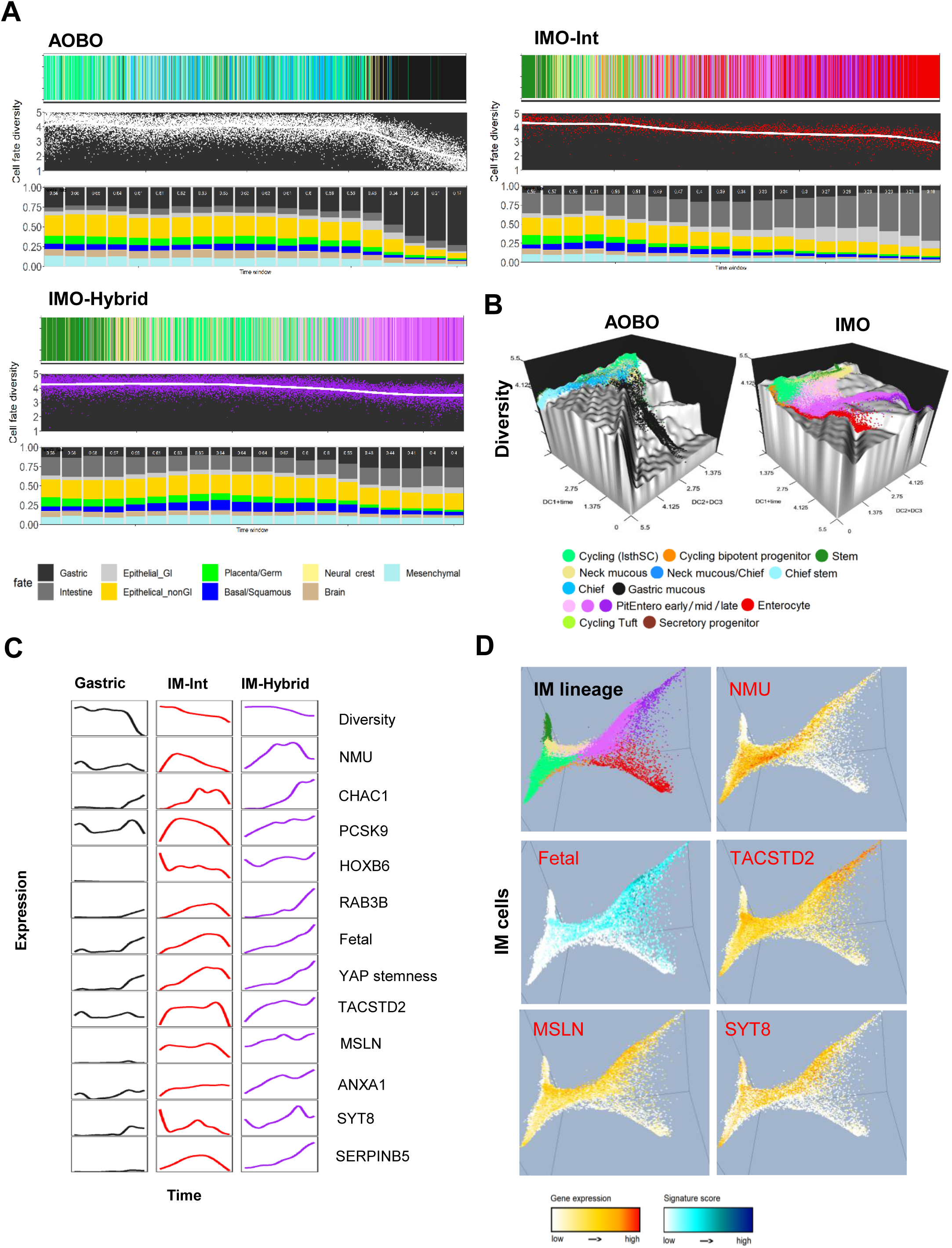
Gastric and IM cell differentiation and cell plasticity/fetal reversion. (A) The AO/BO and IMO trajectories showing the chronological order of cellular differentiation in gastric and IM samples and the alterations in cellular diversity. The x-axis shows the cells ordered by the timeline which was built from the combination of RNAvelocity and diffusion model by CellRank method, without artificial assignment of start/end cells or human-draw arrows of the lineage direction. The IMOs were spit into IMO-Hybrid cells and IMO-Int cells (denoted IM with advanced intestinal differentiation) cells, based on figure 1D. For each organoid group, the top panel shows the cell types of each single cell, as denoted in panel B legend. The second panel shows the cell diversity in the y axis, which gastric lineage is coloured in white, hybrid lineage is coloured in purple and intestinal lineage is coloured in red. The bottom panel shows the predicted cellular proportions of 81 cell fates, then grouped into 9 cell groups coloured according to the legend at the bottom (see supplemental table 7) and averaged at 20 equivalent time windows. (B) Three-dimensional portrayals of the cellular fate diversity states, analogous to mountains, illustrating the dynamic change in cell fates along the differentiation trajectories. Gastric cells proceeded towards a convergent, low-diversity, and predetermined fate, while IM cells maintained their position at the apex of the diversity mountain, akin to the pluripotency observed in stem cells. Within the three-dimensional framework, the x-axis at the base represents the scaled pseudotime combined with DC1, while the y-axis corresponds to the sum of DC1 and DC2. The z-axis, on the other hand, signifies the diversity. (C) The line plot shows the gene expression (y-axis) changes across time trajectory (x-axis) in the gastric, IM-Int, IM-hybrid cells. The diversity score, representative cell fate markers beyond the GI epithelium, fetal and YAP stemness signature scores and representative fetal genes are plotted. (D) 3D diffusion maps of IMOs show the fetal signature score and expression of representative fetal genes (*TACSTD2, MSLN, SYT8*) and basal/squamous marker *NMU* preferentially in the PitEntero cells.

To explore fetal features of the IM cells, we calculated a fetal signature score that was previously found to be elevated in serrated polyps/lesions in the colon,^29^ where the gene list was derived from genes upregulated in murine fetal intestinal (reminiscent of endoderm) organoids compared with adult counterparts^34^ ^35^ (online supplemental table 8). Subsequent studies found high overlap of this fetal signature with genes induced by a yes-associated protein (Yap)-dependent regenerative process of the intestine in response to injury.^36–38^ Therefore, we also derived an intestinal YAP stemness score based on genes induced by Yap in mouse intestinal cells (online supplemental table 8).^36^ The IM-hybrid cells displayed the highest fetal and YAP stemness scores (figure 4C, D), with representative genes such as *TACSTD2*, *MSLN*, *ANXA1*, *SYT8* and *SERPINB5* showing elevated expression, and the highest expression was observed in the PitEntero cell clusters. Overall, in a Wnt-deficient environment, these IM-hybrid cells not only displayed arrested differentiation with lower levels of mature gastric or intestinal markers, but also displayed aberrant differentiation towards other cell lineages and reversion to a fetal and regenerative stem-like state. These features of cellular plasticity and fetal stemness recapitulate those observed in malignant cells, thus providing the mechanistic underpinning of why IM increases the risk of malignant transformation.

Next, we sought to determine whether the scRNAseq data obtained from our organoid differentiation model could accurately represent IM cells in tissues and further explored their relevance to cancer. We first constructed evolutionary clock fitted developmental trees to represent the differentiation trajectories of normal gastric and IM organoid cells (online supplemental methods). In the AO/BO and IMO trees (figure 5A), the terminal nodes represent the observed cell lineage clusters, while the internal nodes reveal hidden ancestor cells during cellular evolution. We mapped an scRNAseq dataset^39^ containing non-neoplastic gastric and cancer tissues to our AO/BO and IMO trees. Epithelial cells derived from the non-neoplastic gastric tissues were separated into normal gastric and IM cell clusters based on their *CLDN3* expression levels. For gastric cancers, we focussed on samples with high *CLDN3* expression, as they were more likely to have evolved through IM. Among the 84 normal gastric cell clusters, 26 were highly correlated (rho.>0.75) with cell nodes of the AO/BO tree (figure 5A, top left), while only 4 clusters were mapped to cell nodes of the IMO tree (figure 5A, top right). In contrast, 14 out of 23 IM clusters were highly correlated with cell nodes of the IMO tree, and none was mapped to the AO/BO tree. These results indicated that our organoid differentiation model can accurately represent the corresponding normal gastric or IM tissues. Normal gastric tissue cell clusters showed high similarities to gastric mucous, chief stem cells and their ancestors, as well as gastric cycling IsthSCs. In addition to similarities to enterocytes, we observed several IM tissue cell clusters with high similarities to PitEntero cells, cycling bipotent progenitors and IM secretory progenitors, suggesting that IM initiation occurs in the stem/progenitor population. We further mapped the cancer cells from 5 IM-type tumours to our AO/BO and IM trees (figure 5A, bottom). Notably, 22 out of 79 cancer cell clusters were strongly correlated with IMO tree nodes, while only 8 out of 79 cancer cell clusters were highly correlated with AO/BO tree nodes. In line with the loss of differentiation in gastric cancer cells, we observed more GC tissue cell clusters with high similarities to IMO cycling IsthSCs, cycling bipotent progenitors and secretory progenitors, where a single GC tended to harbour multiple stem/progenitors. Simultaneously, other tissue cell clusters also co-existed, displaying similarities to enterocytes and PitEntero cells at various stages of hybrid maturation. These results provide strong support for the origination of cancer cells from IM and further highlight the heterogeneity of IM cells arrested at different hybrid states being retained in the cancer cells. In concordance with our observation of increased fetal and Yap stemness signatures in IMOs, we also found a similar increase in IM cells and gastric cancer cells compared with normal gastric cells in tissue samples, further underscoring the role of these signatures in malignant transformation from IM cells (figure 5B).

**Figure 5.**
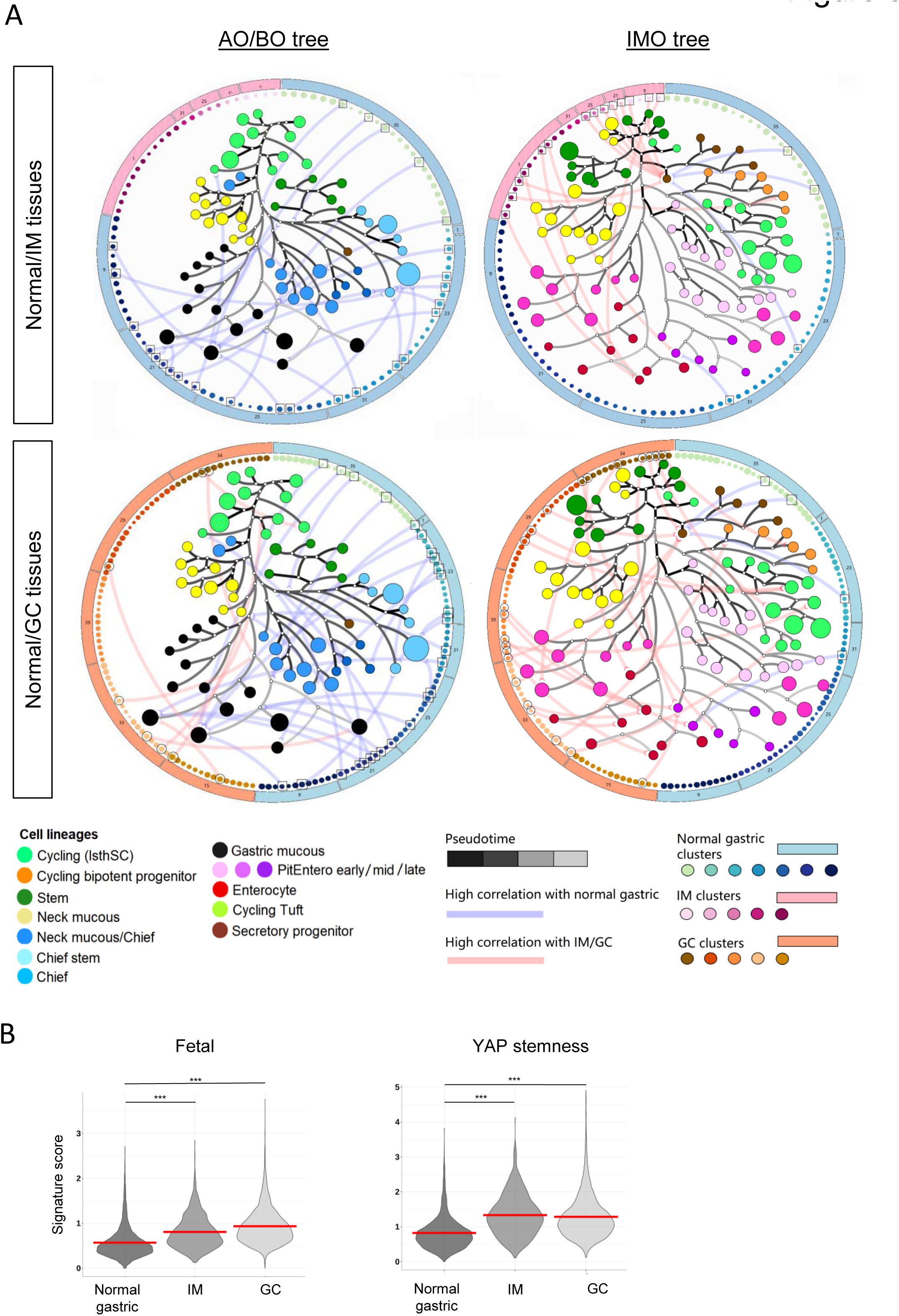
Using developmental trees to compare similarities between human tissue-derived cells to gastric and IM organoids. (A) The lineage tree at the center of each circus plot depicts the evolutionary relationships among gastric (left) and IM (right) organoid cell groups, respectively. Node sizes represent cell numbers, while node colours indicate cell lineages, as shown in the lower left panel. The colour intensity of the tree branches represent the differentiation pseudotime, with darker colours indicating earlier times. Dots at the periphery surrounding the lineage tree represent gastric tissue cell clusters from patient samples, with dot size indicating the number of cells in each cluster and dot colour denoting different patient samples. The outer circle shows the sample ID number for individual gastric samples, coloured correspondingly to normal gastric cells, IM cells, or GC cells as indicated in the legend. Connections denote the top most highly correlated similarities (Spearman correlation R value >0.75) between specific tissue cell clusters and organoid tree nodes, with blue connections representing connections with normal gastric cells, and orange connections representing connections with IM or GC cells. (B) Violin plots showing fetal and YAP stemness scores among epithelial cells derived from normal gastric, IM and GC cells. Red bars indicate the mean for each group. P values were obtained by Mann-Whitney U test, ***p<1×10^-163^.

### IMOs exhibit aberrant methylation of numerous intestinal and gastric genes

Previous studies have demonstrated genome-wide aberrant DNA methylation in IM tissues,^10^ ^13^ ^40^ including up-regulation of several IM genes, such as *CDX1*, *CDX2* and *CDH17*, as a result of promoter demethylation.^14^ ^41^ ^42^ It remains unclear whether epigenetic deregulation may contribute more widely to the IM process beyond the few known genes mentioned above. Principal component analysis using EPIC methylation array data can clearly segregate IMOs from AOs/BOs or colon organoids, indicating that the genome-wide methylation patterns of each organoid type are distinct (figure 6A). Unsupervised hierarchical clustering based on CpG probes showed segregation of the IMOs into two clusters (IMOme cluster 1 and IMOme cluster 2), which were highly concordant with the two IMO gene expression clusters (online supplemental figure 6A). IMOme cluster 1 showed higher concordance with gene expression compared to IMOme cluster 2, and likely had higher IM purity. Next, we integrated both the gene expression and methylation clusters and classified the samples into 12 consensus high purity IMOs and 31 AOs/BOs (online supplemental figure 6A). Notably, the IMOs showed a distinct hypermethylation pattern of CpG islands compared to normal (hyperZ index=2.19, *p*=0.014), as well as a modest level of hypomethylation in the open sea genomic region (hypoZ index=-1.314, *p*=0.094) (online supplemental figure 6A). Subsequently, differential methylation analysis between the consensus IMOs and AOs/BOs clusters identified both hypermethylated and hypomethylated CpG probes (online supplemental figure 6B). By focusing on the anti-correlated methylation and gene expression events, we discovered transcriptional silencing of 456 genes and activation of 453 genes as a result of promoter CpGs methylation (online supplemental figure 6B and online supplemental table 9). When we examined these in relation to the specific genes that we used to define the intestinal and gastric expression signatures, 55 CpGs in 23 genes (including *CLDN18, PGC, GKN1, TFF2* and *TFF1*) from the stomach signature were hypermethylated and down-regulated in IMOs compared to AOs/BOs, and 127 CpGs in 45 genes (including *CDH17, HNF4A, FABP1, MUC2* and *CDX1)* from the colon signature were hypomethylated and up-regulated (figure 6B). These results suggest that aberrant methylation is an important process shaping cell lineage transition towards IM. When comparing our list of differentially methylated IMO genes with a list of previously identified differentially methylated genes in GC,^14^ 23 genes were hypermethylated and down-regulated in both IM and GC, including *CBS*, a gene previously implicated in the CpG Island Methylator Phenotype;^43^ and *PTPN13*, a tumor suppressor frequently mutated in gastric and colorectal cancers.^44^ ^45^ Furthermore, 24 genes were hypomethylated and up-regulated in both IM and GC, including *CDH17*, previously implicated in lymphatic metastasis.^46^ Overall, among the 909 genes deregulated by methylation in IM, 46 are COSMIC genes and 44 of them are known to be mutated in GC.

**Figure 6.**
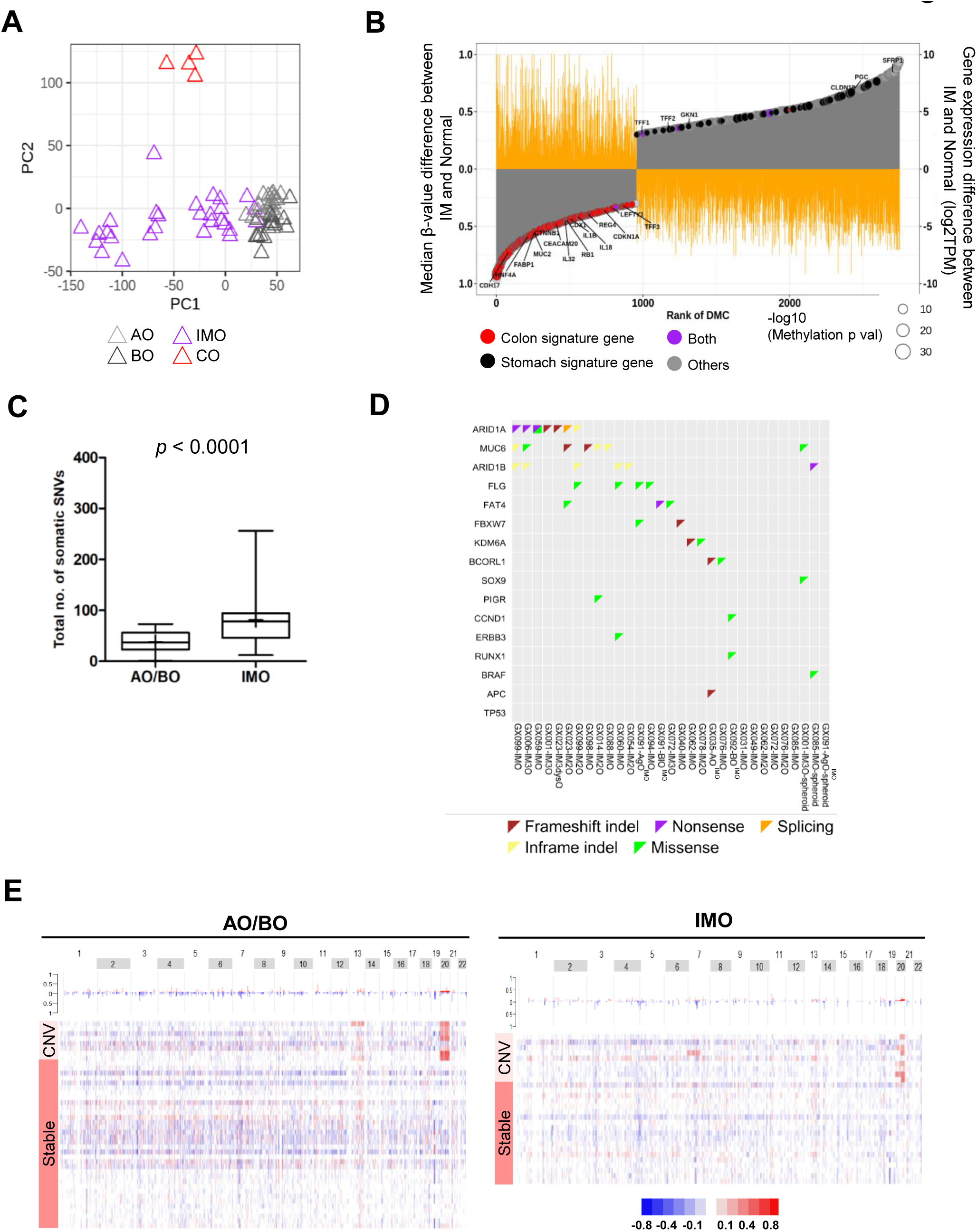
The epigenomic and genomic landscapes of IMOs. (A) PCA plot showing the distinct DNA methylation profiles of each organoid type. (B) Waterfall plot showing the negatively correlated CpG-gene pairs ranked based on the difference in median beta values between IM and normal gastric organoids with corresponding gene expression differences (log2TPM). The colon (red) and stomach (black) signature specific genes, as well as genes found in both signatures (purple) are highlighted. (C) Plot showing the number of SNVs in AOs/BOs *versus* IMOs. The line and the “+” inside the box indicate the median and mean values, respectively. *P*-value was obtained by student’s t-test. (D) Oncoplot showing the mutational profiles of representative IM or tumour drivers in IMOs. Each triangle represents a somatic mutation with the colour indicating the mutation type, as specified in the legend. (E) Heatmap illustrating the chromosomal aberrations in AOs/BOs and IMOs, grouped into chromosomal stable and with CNV groups. (A-E) BO/BfO/BgO/BlO, body organoids; AO/AgO, antrum organoids; CO, intestinal organoids; IMO/IM2O/IM3O/IM3dysO, IM organoids.

### IMOs exhibited elevated mutation rate, but mutations in key cancer drivers were uncommon

Analysis of the WES data showed that the IMOs had an elevated mutation burden compared to the AOs/BOs (figure 6C, D and online supplemental table 10-11), with frequent mutation of *ARID1A (7/28, 25%)* and *MUC6 (6/28, 21%),* less frequent mutation of *APC (1/28, 3.5%)* and *FBXW7 (2/28, 7.1%),* and no *TP53* mutations. Mutation in some other recently reported IM driver genes were also noted. These results are consistent with recent WES studies of IM tissue.^10–12^ While these driver mutations were predominantly observed as single hits, many of them had a minor allele frequency (MAF) well below 0.5, suggesting that they were only present in a subpopulation of IM cells.

We also studied the copy-number variations (CNVs) in the IMOs based on the WES data (figure 6E, online supplemental figure 7 and online supplemental table 12). We observed frequent gain in whole chromosome 20 or 20q, including 7 out of 42 AOs/BOs (16.7%) and 7 out of 28 IMOs (25%). In line with our findings, chr20 gain has been observed in individual IM crypts that were microdissected for WES.^12^ Notably, gain of chromosome 20q is the most frequent type of CNV in GC, and yet our organoid model was able to capture its early emergence in the normal gastric or IM phases. We also observed chromosome 13 gain in GX023-BO and GX094-BfO, chromosome 7 gain in GX035-AO^IMO^, and copy neutral LOH on chromosome 5 in GX023-Imdys3O. The mutational and CNV profiles of our IMOs are distinct from the GCs in the same patient, confirming their non-neoplastic nature and that they are unlikely to be due to tumor contamination (data not shown).

### Progression towards clonal dominance and stability of IM characteristics in long-term IMOs

Next, we examined the effect of long-term culture by randomly selecting 5 IMOs to culture for a continuous period of at least 6 months (IMOLs), followed by WES and RNAseq (online supplemental figure 8). Morphologically, all 5 IMOLs retained their cystic structure (online supplemental figure 8A). Two IMOs, GX001-IM3O and GX076-IMO, which exhibited a low MAF (<0.1) in early passages, showed clonal dominance in long-term culture (online supplemental figure 8B and online supplemental table 11). These results are consistent with what we previously described in normal colon organoids.^21^ The other three IMOs were relatively clonal in early passages and we observed no significant change in their MAF upon long-term culture. At the transcriptomic level, the colon signatures were maintained in most of the IMOs over the long-term and the transcriptomes were stably maintained in long-term culture (online supplemental figure 8C and online supplemental table 3). Overall, despite clonal competition, it was possible to maintain IM characteristics in long-term organoid culture.

### IMOs downregulate FGFR2 and become FGF10 independent

Despite changes in cell plasticity and fetal stemness, there was no statistical difference in the average growth rate between IMOs and AOs/BOs as a group, nor in the pairwise comparison from the same patient (online supplemental figure 9A and online supplemental table 13). Since IM cells showed bi-lineage differentiation potential (figures 1 and 2), we sought to better understand their niche factor requirements. Both AOs/BOs and IMOs died within 2-3 weeks without Wnt-3A, R-spondin1, Noggin or EGF, and within 10-20 weeks without A83-01. However, upon FGF10 withdrawal, all AOs/BOs died within 20-25 weeks in culture, while the COs continued to survive. Moreover, 4 out of 5 IMOs were completely FGF10 independent, as they could be cultured for over 6 months with minimal effects on morphology and growth rates (figure 7A, online supplemental figure 9 and online supplemental table 13). One IMO was partially FGF10 independent, as its growth rate was modestly reduced starting from week 25. To explore the mechanistic underpinnings of FGF10 independence, we examined the expression of FGF ligands and receptors in BOs/AOs and IMOs, and found that *FGFR2*, a cell surface receptor critical for FGF signaling, was expressed in gastric cells in stem and various differentiation stages, while IM cells consistently down-regulated *FGFR2* (figures 2A and figure 7B). Overall, in concordance with the change in cell fate, the growth requirements of the IMOs appeared to shift towards those of intestinal cells, which generally do not require FGF10. We also tested the cellular diversity of the IMOs in FGF10-free conditions and observed an increase in enteroendocrine cell markers, CHGA and CHGB, at both the gene and protein levels (figure 7C). Examination of stem cell markers showed a trend towards increased *LGR5* in the IMOs when cultured in the absence of FGF10 (online supplemental figure 9D).

**Figure 7.**
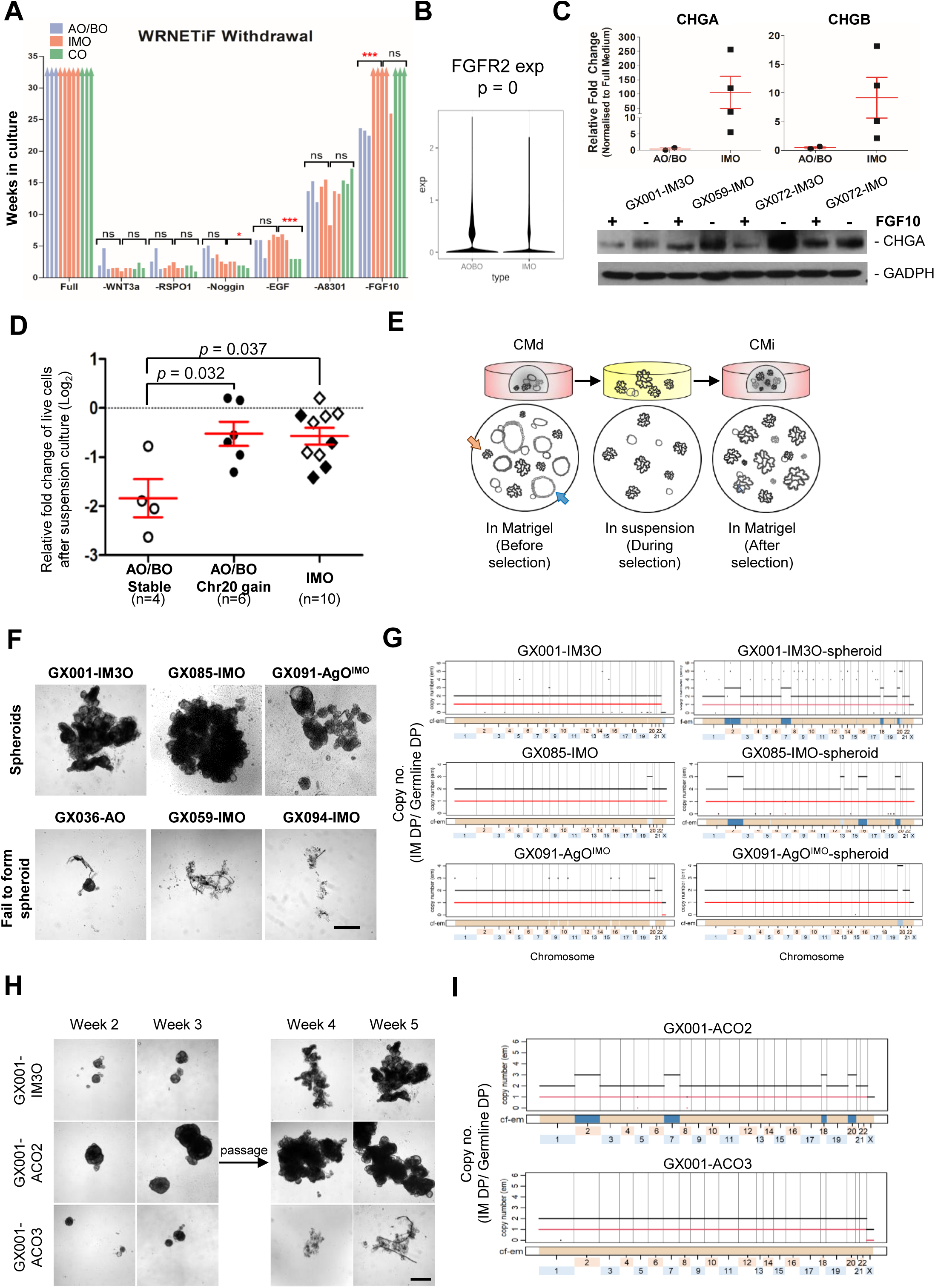
*In vitro* characterization of IM and normal gastric organoids in terms of growth factor dependence and capacity for anchorage independent growth. (A) Bar chart showing the life-span of gastrointestinal organoids under the indicated culture conditions. Bars with arrowheads denote the organoids that can be cultured for more than 6 months. (B) Violin plot showing higher FGFR2 gene expression in AO/BO than IMO. (C) qRT-PCR showing the fold change of *CHGA* and *CHGB* gene expression of organoids grown in a FGF10-free culture medium, relative to those grown in complete culture medium. Error bars represent mean ± SD of 3 technical replicates (top). CHGA protein expression of IMOs grown in full and FGF10-free organoid medium, with GAPDH protein expression as the loading control (bottom). (D) The fold change of live cell number (Log2) between IMOs and AOs/BOs with respect to CNV after 14 days of spheroid selection, relative to initial cell number. Error bars represent mean ± SEM. (E) Schematic diagram showing the process of CMi cell selection by spheroid formation assay. Orange and blue arrows indicate the presence of budding and cystic organoids before the selection process, while budding organoid dominated after selection. (F) Brightfield images showing the 3 IMOs that grown in suspension (upper) and 3 gastric organoids that failed to form spheroids (lower) during selection. Scale bar: 500µm. (G) Copy number plots showing the total (black line) and minor (red line) copy numbers of the corresponding samples, with the estimated cellular fraction profile plotted at the bottom. (H) Brightfield images of GX001-IM3O and its 2 clonal organoids in suspension. GX001-IM3O and GX001-ACO2 were CMi, as they were able to expand in Matrigel-free culture, but GX001-ACO3 was not. Scale bar: 500µm. (I) Copy number plots showing the same CNV patterns between clonal GX001-ACO2 and GX001-IM3O-spheroid (shown in G), while CMd GX001-ACO3 was CNV neutral. Paired student’s t-test was performed between two groups, ns: not significant, ‘*’ *p*< 0.05, ‘***’ *p*<0.001.

### IMOs with chromosomal aberrations displayed a cell-matrix adhesion independent (CMi) growth advantage

We recently showed that almost all GC organoids were cell-matrix adhesion independent (CMi), while normal gastric organoids were not, suggesting that gaining CMi represents the earliest critical step in malignant transformation.^47^ Therefore, we sought to test whether IMOs were CMi by randomly selecting 10 IMOs and 10 AOs/BOs for spheroid formation assay (figure 1A). After 2 weeks in culture, there was a significant growth inhibition, irrespective of the organoid type (figure 7D). However, the IMOs (*p*=0.037), as well as AOs/BOs with chromosome 20 gain (*p*=0.032), were more CMi compared to chromosome stable AOs/BOs. When these 20 organoids were continuously cultured in suspension beyond the initial 2 weeks, most of them died completely by 6 weeks, demonstrating that they were cell-matrix adhesion dependent (CMd). Strikingly, three IMOs (GX001-IM3O, GX085-IMO and GX091-AgO^IMO^) became completely CMi after growing in Matrigel-free conditions for over 3 months (figure 7E-F). Prior to being subjected to spheroid formation assay, GX001-IM3O displayed a mixed cell population with organoids exhibiting a cystic or budding morphology in Matrigel, while GX085-IMO and GX091-AgO^IMO^ were mostly cystic in shape (online supplemental figure 10A). When they were cultured in suspension, a large proportion of organoids died, leaving a small subpopulation that continued to proliferate and eventually became the dominant population. The resulting CMi population of GX001-IM3O consisted primarily of organoids with the budding phenotype, while GX085-IMO and GX091-AgO^IMO^ were still cystic in shape in Matrigel (online supplemental figure 10A). All 3 CMi IMOs remained sensitive to CC dissociation-induced apoptosis and died upon Y-27632 withdrawal (online supplemental figure 10B). As cell-cell adhesion independence (CCi) was previous observed mostly in advanced stage or diffuse type gastric cancer,^47^ their absence in IMOs is consistent with their relatively early stage of neoplastic progression. We did not find any commonly mutated cancer driver genes among the 3 IM spheroids that correlated with the CMi phenotype (figure 6D, online supplemental table 11). Besides, there was low percentage of shared somatic SNVs between organoids and spheroids (online supplemental figure 10C and supplemental table 14). They also did not share any key cancer driver mutations with their paired GC, thereby excluding the possibility of tumor contamination. Intriguingly, we observed more chromosomal aberrations in the IM spheroids compared to their parental organoids (figure 7G and online supplemental figure 10D). All 3 IM spheroids displayed chromosome 20 gain, with evidence of enrichment of this subpopulation from the parental organoids. GX001-IM3O-spheroids showed copy number gains in chromosome 2, 7, 18 and 20 that were not evident in its parental organoid based on WES, but this subpopulation was subsequently detected by scRNAseq and scDNAseq (discussed further below). GX085-IMO-spheroids also showed an extra copy of chromosome 2, 13 and 16. GX091-AgO^IMO^-spheroid gained an additional copy of chromosome 20, increasing from 3 copies in the parental organoid to 4 copies in the spheroid.

To examine the association between CMi and CNVs in IM, clonal organoid lines were derived from GX001-IM3O by manually picking individual organoids, under a microscope based on distinct morphology, for further characterization. The earliest passage of GX001-IM3O was used to generate the clonal organoids to 1) ensure that the presence of CNVs was not an artefact acquired *in vitro*, and 2) to preserve genomic diversity such that clonal organoids with the same chromosomal aberrations as the spheroids could be captured for downstream characterization. Clonal organoids, GX001-ACO2 and ACO3, were subsequently expanded and tested for their CM adhesion independence, along with their parental control, GX001-IM3O. GX001-ACO2 formed a compacted cell cluster and was completely CMi, while GX001-ACO3 failed to propagate and died at week 4 (figure 7H). The two clonal IMOs showed no differences in their growth rate or their niche factor requirements compared to the parental GX001-IMO (online supplemental figure 11A-B). GX001-ACO2 shared the same CNV pattern as GX001-IM3O-spheroid, with copy number gains in chromosomes 2, 7, 18, and 20, while GX001-ACO3 was chromosome stable (figure 7I).

### IMOs with chromosome 20 gain were associated with CM adhesion independence, and IM spheroids displayed stemness and tumor-like features

To better understand the significance of chromosome 20q gain in the 3 IM spheroids, we subjected them to scRNAseq analysis and compared them to their parental IMOs. We inferred the CNV status of individual cells and found that gain of chromosomes 2, 7, 18 and 20 in GX001-IM3O-spheroid was indeed enriched from a subpopulation that pre-existed in its parental organoids (figure 8A). The presence of this minor cell cluster in GX001-IM3O was further validated at the DNA level by single-cell CNV (scCNV) analysis (online supplemental figure 11C). Similarly, gain of chromosome 2 in GX085-IMO-spheroid was enriched from a small fraction of cells in GX085-IMO. Chromosome 20 gain was found in GX091-AgO^IMO^, and subpopulations of GX091-AgO^IMO^-spheroid showed additional gain of chromosomes 2, 5, 8 and 16, and loss of chromosome 4, along with an increase in the inferred CNV score for chromosome 20 (figure 8A), which was consistent with the WES data showing that the spheroid population actually gained one more copy of chromosome 20 to become 4N (figure 7G). Moreover, we inferred the single-cell CNV status of other gastric and IM organoids, and confirmed the presence of chromosome 20 gain in 2 normal organoids, GX023-BO and GX036-AO, both of which were CMd (figure 1A and online supplemental figure 12). Thus, we hypothesized that both IM and chromosome 20 gain may be needed to set the stage for CMi growth.

**Figure 8.**
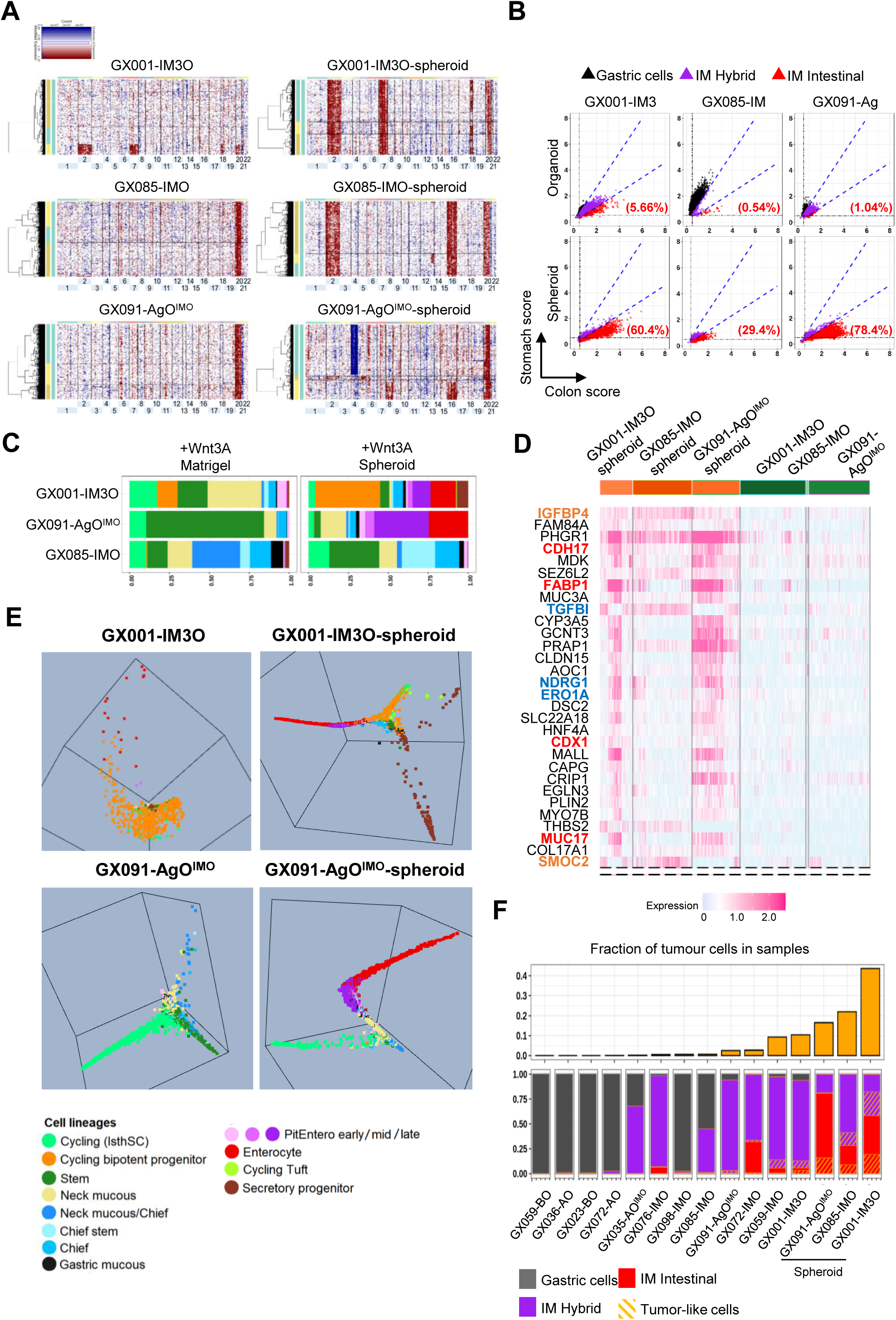
Single-cell transcriptomic analysis comparing paired IM spheroids and organoids. (A) Hierarchical clustering of single cells (rows) from 3 IM organoid-spheroid pairs based on inferred copy number from scRNAseq data. The cells with copy number gain and loss are indicated by red and blue colour, respectively. (B) Scatter plots showing the cellular composition among the IM organoid-spheroid pairs (percentages indicate intestinal lineage cells). (C) Stacked bar charts showing the cell type composition of each sample when cultured in Matrigel (left) or as spheroid (right). Each cell type is coloured according to the legend in (E). (D) Heatmap showing the top 30 upregulated DEGs among 3 IM spheroid-organoid pairs using a non-parametric Wilcox rank sum test. Stem cell markers are labeled in orange, intestinal lineage markers are labeled in red and hypoxia-induced genes are marked in blue. (E) A three-dimensional cell lineage trajectory plot, through diffusion map analysis, showing the adoption of different differentiating cell lineages in GX001-IM3O-spheroid and GX091-AgO^IMO^-spheroid, compared with their parental organoids (only the subpopulation with chromosome 20 gain is displayed in GX001-IM3O for comparison with its corresponding spheroid). (F) Bar chart (top) showing the proportion of tumour-like cells identified by Scissor algorithm in organoid samples in ascending order. Bar chart (bottom) showing the proportion of gastric, IM-hybrid and IM-intestinal cells in each sample, indicated by colour, with the fraction of tumour-like cells highlighted by diagonal orange lines.

Next, using stomach and colon signature scores, we identified a substantial enrichment of IM cell fractions with increased colon scores in the spheroid samples and thus, a tendency to adopt a more differentiated phenotype (figure 8B and online supplemental table 5). When cultured in Matrigel, the IMOs (GX001-IM3O and GX091-AgO^IMO^) showed enrichment of stem and cycling IsthSCs or bipotent progenitors (figure 8C, E). However, when they were cultured in suspension, they displayed a wide spectrum of differentiation, ranging from stem cells, cycling bipotent progenitors, secretory progenitors, as well as higher proportions of PitEntero and enterocyte cells, despite the presence of Wnt-3A and other stem cell enriching niche factors. Moreover, in Matrigel-free conditions, the gastric cell subpopulation that was originally found in GX085-IMO was completely replaced by hybrid and intestinal cells (figure 8B).

We performed differential gene expression analysis, with the gastric cells filtered out, on the 3 IM-spheroid/parental organoid pairs to identify key transcriptomic changes that could contribute to CMi in IM cells (figure 8D). Interestingly we noted concurrent up-regulation of intestinal stem markers (e.g. *SMOC2*, *IGFBP4*), as well as intestinal differentiation markers (e.g. *FABP1*, *CDH17*) in the IM-spheroid samples, suggesting an enriched cell diversity (figure 8C, E). Other up-regulated genes included *THBS2* and *MDK*, which are oncogenic proteins that can serve as ligands for integrin alpha 6 and beta 1,^48^ ^49^ as well as *NDRG1, ERO1A* and *TGFBI*, which are known hypoxia-induced genes that play a pivotal role in the EMT process and promote tumor metastasis.^50–52^

Lastly, we used Single-Cell Identification of Subpopulations with Bulk Sample Phenotype Correlation (Scissor)^53^ to detect whether the IMOs contain subpopulations of cells with cancer phenotypes. Briefly, an expression matrix from bulk RNAseq data from 38 normal and 43 gastric cancer organoids, as well as scRNAseq data from 4 normal gastric organoids, 8 IM organoids and 3 IM spheroids, was used to build a correlation matrix and a cell-cell similarity network. A phenotype of either ‘normal’ or ‘tumor’ was then integrated into the network regression model to check for tumor-like cells (figure 8F). The Scissor method showed less than 1% of cells were tumor-like in all AOs/BOs and in 4 IMOs (3 without chromosome 20 gain). Strikingly, it detected ∼2-10% of IM cells as tumor-like in 4 IMOs: GX091-AgO^IMO^ and GX001-IM3O, which carried chromosome 20 gain, and GX072-IMO and GX059-IMO, which were derived from advanced IM tissue (figure 8H). More importantly, the 3 IM spheroid samples, which were composed of either hybrid and/or intestinal cells, contained 16-40% of tumor-like cells. Overall, we demonstrated that IM cells (hybrid or intestinal) with chromosome 20 gain were highly associated with a cancer phenotype in addition to being CMi.

## DISCUSSION

Here, we present the establishment and comprehensive multi-omics characterization of the largest collection of IMOs (n=28) from human IM tissues to date. Since the IMOs were established from resected tissue specimens covering a large area of the mucosa, we were able to capture a diverse array of the glands that harbor heterogeneous level of genetic and epigenetic perturbations, and along with them the stem or progenitor cells with divergent differentiation potential and anchorage independent ability in culture. Specifically, we noted organoid models displaying the full spectrum from hybrid to advanced intestinal differentiation, which can serve as cell models to reflect the underlying biology of the IM process beyond the previous histologically defined incomplete or complete intestinal metaplasia.

The scRNAseq data from our gastric and IM organoid models, which are enriched in immature components, complement data derived from human gastric tissue by revealing the detailed subtypes and developmental trajectories of the various stem and progenitor populations. This high-resolution data enabled us to reveal sub-populations of IM cells that displayed similarities to murine intestinal stem cell subgroups under homeostasis or injury-induced regeneration that were revealed by lineage tracing experiments. Previous studies in mice have identified different subpopulations of colon cells beyond the Lgr5+ stem cells, such as the transit amplifying bipotent progenitors, the secretory or enteroendocrine progenitors and the early enterocytes, which can dedifferentiate into ISCs and regenerate the entire intestinal crypt upon various types of injuries.^54–57^ Currently, there is intense interest in studying the relevance of these different stem cell subtypes to serve as the cancer initiating cell. Our IMO secretory lineage cells expressing *PROX1*, *BMI1* along with *ASCL2* and *MEX3A*, highly re-capitulate the mouse Prox1+ cells, which has been shown by lineage tracing to be able to generate tuff and EE cells, and can also act as reserve stem cells to regenerate the whole crypt upon injury.^28^ Interestingly, in colorectal cancer, *MEX3A* marked a population of drug-resistant cancer stem cells that can transiently adopt a YAP+ fetal progenitor state upon chemotherapy.^58^ Concordantly, recent studies have identified a YAP-dependent regenerative process that do not contribute to homeostasis in the intestine, but is important for regeneration upon various types of injury induced by infection, inflammation or irradiation.^36–38^ These regenerative stem cells showed fetal lineage reversion, reduced dependency on Wnt, and upon oncogenic mutation, could become cancer stem cells.^59^ Our hybrid IMOs, which displayed high expression of fetal and YAP signatures especially when grown in Wnt deficient conditions, may mimic these regenerative stem states. Furthermore, a recent study showed that co-culture of patient metaplastic fibroblasts or cancer-associated fibroblasts with gastroids can induce proliferation and *TACSTD2* (also known as *TROP2*) expression and promote dysplastic progression. While the detailed transcriptomic profile of the *TROP2+* gastroid cells is not known, they may represent fetal reversion induced by stromal factors.^16^ Interestingly, a mouse organoid model of serrated neoplasia pathway harboring RSPO fusion/*KRAS*/*TP53*/*SMAD4* quadruple mutations developed fetal lineage reversion concurrent with gain of Wnt independence upon transient exposure to TGFβ.^60^ Thus, the adoption of a fetal intestinal phenotype in the hybrid IMOs may facilitate growth adaptation when niche factors are deficient. Concordantly, in a recent study of metastatic colon cancer, transition through a fetal state to aberrant differentiation foci was noted in some liver metastasis and these phenotypes could be re-capitulated in metastatic colon cancer organoids when niche factors were withdrawn, which was highly similar to our observations in IM cells.^31^ A previous study on a collection of colonic polyps revealed that gastric metaplasia and reversion to a fetal phenotype is an important cancer pathway in serrated neoplasm.^29^ Here, we showed that intestinal metaplasia of gastric cells also displayed a similar reversion to fetal stemness that could contribute to cancer progression. Since lineage plasticity confers adaptation to different environmental stresses and can facilitate drug resistance and survival in metastatic distant organs, their emergence in the IM process in stomach, as well as the serrated pathway in colon, could serve to explain the relatively more aggressive nature of gastric cancer or serrated colon cancer compared with conventional colon cancer. Overall, neoplastic progression of the gastrointestinal tract seems to share common features through the metaplastic process, with concurrent reactivation of the fetal phenotype and lineage plasticity.

Consistent with previous studies that showed similarities between early gastric cancer and the IM stem lineage,^11^ ^19^ our study further identified subpopulations within gastric cancer that exhibit expression profiles similar to various IMO subpopulations, including secretory progenitors, cycling bipotent progenitors, cycling isthmus stem cells and PitEntero cells. It would be interesting to investigate whether these different IM stem characteristics in cancerous tissue may confer differential resistance to chemotherapy or irradiation. We further confirmed a graded elevated fetal signature from IM to gastric cancer *in vivo.* Indeed, fetal gene such as *TACSTD2*, also known as Trop2, has been demonstrated to be expressed in human incomplete intestinal metaplasia, all dysplasia and majority of gastric cancer.^61^ Another fetal gene *MSLN* is aberrantly expressed in many solid cancers and constitutes a cancer antigen target for CAR-T therapy.^62^ Future studies to exploit fetal reversion or lineage plasticity as therapeutic options for IM and GC are plausible.

We also showed that during the metaplastic process, the growth requirement of gastric cells changed to become similar to those of colon cells, with downregulation of *FGFR2* and reduced reliance on the availability of FGF10. Recent studies in gastric and Barrett’s mucosal tissue identified distinct fibroblast subsets in IM.^11^ ^16–18^ It is unclear whether changes in IM epithelial cells initiate stromal remodeling or vice versa. Therefore, it would be interesting to examine this epithelial-stromal interaction *in vitro* using our IM organoid models and corresponding mucosal fibroblasts, enabling functional dissection of cellular interactions in these unique cell populations and potentially identifying blocking mechanism to reverse IM.

Our study also revealed the neoplastic potential of the IM process, with the emergence of cells carrying CNV and anchorage-independent growth ability. In a previous study, long-term patient follow-up showed that somatic copy number alterations (sCNAs) were associated with progression towards dysplasia or early GC.^10^ Our observation of frequent chromosome 20 gain in up to 25% of IMOs, though not observed in this previous study, has been confirmed in another study through microdissection of individual IM crypts or clusters of glands in the stomach.^12^ Interestingly, this study found that IM glands with chromosome 20 gain tend to expand to colonize the field, indicating a growth advantage *in vivo*, thus supporting our observation that chromosome 20 gain in IMO favours CMi. Interestingly, one adaptation manifested by these CMi cells is the adoption of enhanced cell diversity that featured a mixed population of stem (*SMOC2* high) and differentiated cells (*FABP1* high). This highly re-capitulate a previously described colorectal cancer liver metastases mouse model, in which the initial undifferentiated cells in micrometastases had to segregate into stem and differentiated compartments before metastatic colonization could expand to form macrometastases.^63^ While CNV commonly occurs in cancer subsequent to *TP53* mutation, we found that in the stomach, CNV in forms of broad chromosome gain can precede *TP53* mutation. Despite the absence of key cancer driver mutations, these IM cells with chromosome 20 gain were able to display cancer phenotypes based on their transcriptomes, as depicted by our Scissor analysis, with expression of oncoproteins, such as MDK and the ability to gain anchorage independent growth. Given that expression of the IM transcriptome and chromosome 20 gain are very common in GC,^14^ ^15^ it is plausible to hypothesize that these cells may represent the earliest neoplastic precursor of IM-induced GC. Overall, our study highlights a unique model of carcinogenesis that is secondary to chronic inflammation, characterized by epigenetic perturbations leading to the metaplastic process that triggers lineage plasticity and fetal lineage reversion, coupled with CNV and the unique adaptation of enforced stem and differentiation compartments to achieve an anchorage independent phenotype. Our study is limited by the relatively small number of samples displaying CMi, and detailed follow-up mechanistic studies are needed to understand how together, CNV and IM can cause CMi. Nonetheless, the IMO models established here, capturing different injury repair progenitors that display lineage plasticity, offer an ideal platform for studying the biology of early stages of cancer evolution, and for evaluating potential therapeutic opportunities in cancer prevention and treatment through targeting IM and their corresponding plasticity.

## Data Repository

The WES, methylation array and RNAseq data will be deposited into the European Genome-Phenome Archive. Normalized scRNAseq, and scCNV data will be deposited to GEO.

## Acknowledgements

We thank clinicians in the Hong Kong West Cluster, Hospital Authority for clinical care and sample coordination. We thank the Genomics and Bioinformatics Cores of the Centre for PanorOmic Sciences (CPOS), The University of Hong Kong for performing WES, EPIC methylation array, RNA-seq and single-cell sequencing experiments. We thank the Imaging and Flow Cytometry Core of the CPOS for providing the single cell sorting, flow cytometry and imaging technical supports. This work was substantially supported by a theme-based research grant (Project No. T12-710/16-R) from the Research Grants Council of the Hong Kong SAR, China; and partly supported by the Centre for Oncology and Immunology under the Health@InnoHK Initiative funded by the Innovation and Technology Commission, The Government of Hong Kong SAR, China; a donation from the Hong Kong Jockey Club and the Kadoorie Charitable Foundation. Functional characterization of organoids were partially supported by the General Research Fund (Project No. 17108616) and scRNAseq of intestinal organoid by General Research Fund (Project No. 17107219) from the Research Grants Council of the Hong Kong SAR, China.

## Author Contributions

HHNY, SYL conceived, designed, and supervised the study. SSKY, SLH, WYT, DC, ASYC, SWY, HSH, STY, SYL, HHNY collected data, performed experiments and/or analyzed data. YT and HCS performed the bioinformatics data analysis. SYKL contributed clinical data and samples. YH provided advice on bioinformatic analysis of scRNAseq data. HC contributed protocols and/or reagents and critical comments on the manuscript. SYL, HHNY, SSKY and YT wrote the manuscript with assistance from ASC. All authors contributed comments and approval for the manuscript.

## Declaration of Interests

SYL and STY have received research sponsorships from Pfizer, Merck, Servier. HC is inventor on patents related to organoid research. His full disclosure: www.uu.nl/staff/JCClevers/Additional functions. The other authors declare no competing interests.

## Ethics Statement

This study involves human participants and was approved by the Institutional Review Board of the University of Hong Kong and the Hospital Authority Hong Kong West Cluster (IRB reference ID: UW14-257).

## Online Supplemental Methods

### Patient cohort and clinical specimens

Fresh human tissues were collected from patients who underwent gastrectomy for gastric cancer or colectomy for colorectal cancer at Queen Mary Hospital, Hong Kong. All patients signed informed written consent, and the research protocols were approved by the Institutional Review Board (IRB) of The University of Hong Kong and the Hospital Authority Hong Kong West Cluster. Gastric mucosa tissues, including intestinal metaplasia (IM) and normal stomach mucosae were sampled from gastric antrum or body regions away from the tumour, and dissected from underlying muscularis propria by pathologists. Mucosal strips (∼1cm x 0.8cm in area) were bisected, with half of the tissue snap frozen and the other half used for organoid culture. Cryostat sections were examined by a pathologist to classify the mucosa as normal body, normal antrum, or IM. The presence of IM was graded on a scale from 0 to 3, based on the degree of gastric intestinal metaplasia observed per hematoxylin and eosin (H&E) stained section. IM0 indicates 0%, IM1 indicates ≤30%, IM2 indicates 30-70%, and IM3 indicates >70% of the area was affected by IM within a specific mucosal tissue sample. Normal colon or ileal mucosa were dissected from the underlying muscle and submitted for frozen section and organoid culture, serving as a reference for comparison.

### Patient and Public Involvement statement

Patients or the public WERE NOT involved in the design, or conduct, or reporting, or dissemination plans of our research

### Establishment of human organoids from fresh tissue

Gastric glands from IM and normal tissues were isolated for organoid culture using previously described methods.^1–4^ Briefly, fresh tissues were rinsed with 1x PBS buffer containing 100µg/mL Primocin and 1x Penicillin/Streptomycin (P/S). Stromal and fatty tissues were removed from the specimens using scissors, and the mucosa was cut into small pieces (∼0.5cm x 0.5cm), followed by incubation in a PBS-based chelation buffer (10mM EDTA, 0.5mM DL-dithiothreitol, 1x P/S, 10µM Y-27632 and 100µg/mL Primocin) at 4°C for 1 hour, with shaking. After incubation, the tissue pieces were placed on a petri dish and mechanically pressed with a glass-slide to release the gastric glands, followed by transfer to a clean 50mL falcon tube with 1xPBS and shaken vigorously for 1-2 minutes. The released glands were collected by centrifugation at 200-400 x g for 5 minutes, washed twice with 1xPBS and once with Advanced DMEM/F12. Afterwards, the cell pellets were re-suspended in Matrigel and 50µL of the cell suspension was seeded in each well of a pre-warmed 24-well plate. Both normal gastric and IM organoids were cultured using a standard human gastric organoid culture medium (advanced DMEM/F12 supplemented with 1x P/S, 1x HEPES and 1x Glutamax, 50ng/mL EGF, 10% Noggin conditioned medium, 10% Rspo1 conditioned medium, 50% Wnt-3A conditioned medium, 100ng/mL of FGF10, 1nM of Gastrin, 2µM of TGFβi (A83-01), 1x B27 and 1mM N-Acetylcysteine). Seven normal colon organoids and 3 organoids derived from normal ileum, described in our previous study,^5^ were used as references in this study. They were cultured using a standard human colon organoid culture medium, as previously described.

For routine passaging, normal stomach, intestinal and IM organoids were mechanically split using fire-polished glass Pasteur pipettes, washed with Advanced DMEM/F12 and embedded in Matrigel at a 1:4 ratio. After each passage, 10µM Y-27632 was added to culture medium to suppress cell dissociation-induced apoptosis. Five IM organoids were cultured continuously for over 6 months and passaged once weekly. All cultures were regularly checked and confirmed to be mycoplasma free.

### DNA/RNA extraction from human organoids

Six to eight Matrigel droplets containing organoids at 90% confluence were incubated with Cell Recovery solution (BD bioscience) at 4°C for 30 minutes and washed twice with PBS. The organoids were then lysed in 1.2mL RLT lysate buffer provided by the AllPrep® DNA/RNA/miRNA Universal Kit (Qiagen). DNA and RNA were extracted from the organoids following the manufacturer’s protocol. For qRT-PCR, total RNA was extracted from the organoids using the miRNeasy Mini Kit (Qiagen) according to the manufacturer’s protocol.

### Histological and immunohistochemical analysis of human tissues and organoids

Human biopsies were frozen in liquid nitrogen and embedded in OCT compound prior to cryostat sectioning. To generate cell blocks that could be sectioned, organoids from 4-6 Matrigel droplets were incubated with Cell Recovery Solution, as described above. The released organoids were re-embedded in 50µL of Matrigel, seeded into a pre-cast 3% agarose gel mold and allowed to solidify at 37°C for 15 minutes. The cell blocks were gently released from the molds with a toothpick and fixed in neutral buffered formalin (pH7.0) overnight, and then embedded in paraffin. Both the frozen tissue blocks and organoid paraffin blocks were sectioned at 5-6µm thickness and subjected to H&E staining for pathological assessment, as previously described.^4^ Immunohistochemistry (IHC) of CDX1 and CLDN18 was performed on organoid sections using the Roche Ventana Discovery Ultra automatic slide stainer after heat-mediated antigen retrieval. Selected normal gastric and normal intestinal organoids were included as biological controls. The organoid sections were incubated with rabbit anti-human CDX1 (1:1000, Invitrogen) or mouse anti-human CLDN18 (1:1500, Invitrogen) antibodies at 4°C overnight, and the IHC reaction was detected using the OptiView DAB IHC Detection Kit (Ventana), according to manufacturer’s instructions. The H&E and IHC images were captured using a Nanozoomer digital slide scanner system (S210, Hamamatsu Photonics) and viewed using NDP.view2 (Version 2.8.24). Bright-field microscopy images of the organoids were captured using a Color Imaging Microscope (Axio Observer 7, Zeiss) with a high-speed CMOS colour camera (Olympus DP74) installed.

### Transcriptome profiling and data analysis

RNA sequencing (RNAseq) was performed on 80 organoids (28 IM, 42 normal gastric and 10 normal intestinal organoids), 59 frozen tissues (25 IM, 27 normal gastric and 7 normal intestinal tissues), and 5 IM organoids in long-term culture (IMOLs) (online supplemental table 2). Four micrograms of total RNA per sample were used for cDNA library construction using the KAPA Stranded mRNA-Seq Kit (KR0960-v3.15), according to the manufacturer’s protocol, and were sequenced as 151bp paired-end reads using the NovaSeq6000/HiSeqX Ten platform, or 101bp paired-end reads using the HiSeq1500. The RNAseq reads in FASTA format were aligned to the Genome Reference Consortium human build 37 (hg19) by STAR (version 2.5.2b).^6^ Duplicate reads were removed from the BAM files and transcript abundance was quantified by RSEM^7^ to calculate the transcripts per million (TPM) value for each gene.

Unsupervised hierarchical clustering was performed on the organoid cohort using the mean centered log2 TPM values by Cluster 3.0 (Version 1.58).^8^ Genes with low expression (maximum gene expression in log2 TPM < 0) or low variation of gene expression (VAR<1) across all samples were removed before the average linkage clustering was performed. The results were visualized as a heatmap using Java TreeView 3.^9^ We used protein expression data from The Human Protein Atlas (HPA) (version 17) to derive stomach and colon gene signature scores for each sample in our cohort. The HPA provided immunohistochemical data demonstrating the unique or high expression of specific proteins in a particular organ compared with other tissue types.^10^ ^11^ We then utilized this protein-based information to create organ-specific gene lists and subsequently calculated the organ-specific gene signature scores using our transcriptome data. The stomach and colon signature scores of individual samples were calculated by averaging the log2 TPM values of the genes in each organ-specific list. Additionally, we calculated an aggregate score for each samples by first multiplying the log2 TPM values of stomach signature genes by -1, and then subsequently averaging these values together with the log2 TPM values of the colon signature genes.

### Single cell RNA sequencing and data analysis

Single cell RNA sequencing (scRNAseq) was performed on 8 IM organoids, 4 normal gastric organoids, 3 normal intestinal organoids, 3 IM spheroids and 1 IM fresh tissue, (online supplemental table 4), with the organoids cultured both with or without Wnt3A culture medium for 4 days to promote cell differentiation. Single cell suspensions were prepared as previously described.^5^ Briefly, organoids were trypsinized into single cells and stained with 10µg/ml DAPI (Thermo Fisher Scientific) in a PBS based staining buffer (2% FBS, 1xB27 and 10µM Y-27632) for 2 minutes at room temperature in the dark. The single cells were washed, centrifuged at 1,000 x g for 5 minutes and re-suspended in the staining buffer at a cell concentration of 1×10^6^ cells/mL for sample sorting. The cell suspension was passed through a 40µm cell strainer before performing FACS. Live single cells were gated using standard parameters for forward verses side scatter areas, excluding dead cells with positive DAPI signals, and collected in 1xPBS with 0.04%BSA at a final cell concentration of 1,000-2,000 single cells/µL for cell encapsulation. Single cell barcoding, reverse transcription (RT), cDNA amplification and scRNA-seq library construction were performed using a Chromium Single Cell 5’ Library and Gel Bead Kit (10X Genomics, 16 rxns PN-1000006), according to the manufacturer’s instructions. The constructed libraries were sequenced using a NovaSeq 6000 platform (Illumina) with read lengths PE151. Afterwards, the sequencing reads were aligned to the human genome (GRCh38) using CellRanger (Version 2.2.1). Unique molecular identifier (UMI) count matrices were pre-processed and filtered by scatter R package (version 1.10.0, BioConductor).^12^ Single cells with very low or high UMI counts were removed, as were cells with a high percentage of mitochondrial (Z-score > 2) or ribosomal reads (Z-score > 2), low expression of housekeeping genes (*GAPDH* & *ACTB*), as well as cells that did not show correlation between gene counts and read counts using a robust linear model in MASS (R package, Version 7.3). Furthermore, genes with expression in less than 10 total counts across a dataset or with expression in less than 5 individual cells were removed. The “LogNormalize” method in Seurat (R package, Version 3.2.3) was used to normalise the single-cell gene expression data and the doublets were removed by DoubletFinder (R package, Version 2.0.2). The final dataset comprised of 92,730 organoid cells (online supplemental table 4).

The stomach and colon signature scores for single cells were calculated by the Single Cell Signature Scorer^13^ using the same stomach and colon gene sets as described above. We incorporated our previously published intestinal organoid dataset encompassing two colon samples, and one new ileal sample cultured with or without Wnt3A as reference to derive the signature scores of normal intestinal cells.^5^ The cell subgroups, namely gastric, hybrid, and intestinal cells, were defined using GX036-AO and H019-NO as references to draw the boundary of gastric or intestinal cells respectively. We defined gastric cells as those with a colon score to stomach score ratio of ≤ 0.75; hybrid cells as those with a colon score to stomach score ratio between 0.75 and 1.75, intestinal cells as those with a colon score to stomach score ratio ≥ 1.75.

To examine cell cycle variation in our data, the CellCycleScoring function in Seurat (R package, Version 3.2.3) was used to assign each single cell a cell cycle score based on their expression of G2/M and S phase markers. Thus, each cell was predicted to be in G2/M or S phase, and cells that did not express either G2/M or S genes were assumed to be non-cycling and in the G1 phase.

### Construction of gastric/IM cell lineages at single-cell level

A gastric cell atlas was constructed using the scRNAseq data from 15 IM organoids, 8 normal gastric organoids and 3 IM spheroids. The data was integrated and the batch effects were removed by the “multiBatchNorm” and “batchCorrect” functions in Batchelor (R package, version 1.4).^14^ The batch-corrected dataset was scaled and clustered by “FindClusters” in Seurat (version 3.2.3, R package)^15^ after regressing out the cell-cycle differences between the S phase and G2M phase. 36 cell clusters (C0 to C35) were identified with distinct gene expression patterns (cell resolution=2) (online supplemental figure 4A).

Since the organoid cultured cell clusters primarily consist of relatively immature cells, including stem cells, progenitors and precursors of various specific lineages at different levels of differentiation, traditional cell mapping methods such as human cell landscape (HCL)^16^ analysis are not applicable. Instead, we used known cell lineage markers that characterize various developmental stages of gastric and intestinal cells (online supplemental table 6), and calculated the average expression (Z-score) of specific lineage marker genes after removing non-expressed genes (defined as expressed in <20% of cells) in each cluster. Cell clusters were then grouped into 14 different cell types based on the relative expression of these lineage marker genes (online supplemental figure 4B, C). Five clusters were grouped as “Stem” due to their high Z scores for stem cell genes (*ASCL2*, *OLFM4*, *LGR5*, *SMOC2*, *EPHB2*). Six clusters were grouped as cycling cells because of their high expression of proliferation cell markers (*BIRC5*, *MKI67*, *PCNA*, *TOP2A*, *HELLS*) and because a majority (>60%) of the cells were predicted to be in S or G2/M phase by their cell cycle score. Since they also expressed high levels of a previously reported gastric isthmus stem cell marker *STMN1*,^17^ we refer to this group as “cycling isthmus stem cells” (IsthSC). Another cluster of actively cycling cells (Cluster C21) was only observed in IMO samples, expressing both enterocyte (*FABP1*, *KRT20*) and goblet cell markers (*HES6*, *SPINK4*), along with proliferation genes, mimicking bipotent progenitors in the intestine. We refer to this group as the “Cycling bipotent progenitors”. We grouped 4 clusters as “neck mucous” cells and 3 clusters as “chief” cells due to their high expression of neck mucous (*MUC6*) or chief cell (*PGC*) markers, respectively. A separate cluster with co-expression of *MUC6* and *PGC* is referred to as “neck mucous/chief” cells. Another cluster showed high expression of both chief cell marker *PGC* and stem markers (*LGR5*, *TNFRSF19*), and we refer to them as “chief stem” cells. Three clusters showed the highest Z scores for *TFF2*, *TFF1* and *MUC5AC*, genes expressed in the pit or antral mucous glands, and as a result, we refer to them as “gastric mucous” cells. Five clusters expressed both gastric mucous markers (*TFF2*, *TFF1*, *MUC5AC*) and enterocyte markers (*FABP1*, *KRT20*, *ANPEP*), with high Z scores. We refer to these as “PitEntero” cells, which was consistent with each single cell showing bi-lineage differentiation towards gastric pit mucous cells, as well as enterocytes. To indicate the progressive stages of IM, we further subclassified them as “early”, “mid” or “late”, based on the progressive increase in mature enterocyte markers *KRT20* and *ANPEP*, and a corresponding decrease in the gastric pit marker *MUC5AC*. One cluster, despite persistent expression of gastric lineage markers such as *TFF2* and *ANXA10*, showed the highest expression of all 3 enterocyte markers (*FABP1*, *KRT20*, *ANPEP*) and the lowest expression of *MUC5AC*. We refer to these as “enterocyte” cells and consider them to be representative of the most advanced IM cells. Thus, our “PitEntero” and “Enterocyte” classifications effectively capture the progressive stages and levels of IM transformation at the single-cell level.

Two cell clusters expressed high levels of secretory, enteroendocrine or tuft cell markers. One of these clusters was actively cycling (>80% cells in S/G2M phase), and displayed high expression of tuft cell markers (*AVIL*, *POU2F3*), leading us to refer to them as “cycling tuft” cells. In contrast, the other cluster was quiescent and expressed multiple secretory lineage markers, including tuft (*AVIL*, *POU2F3*), enteroendocrine (*NEUROD3*, *SYP*, *CHGA*), and goblet (*HES6*, *SPINK4*) markers. Moreover, this cluster expressed enteroendocrine progenitor markers (*PROX1*, *BMI1*) and the injury inducible stem marker *MEX3A*, along with high expression of the stem marker *ASCL2*. We refer to this group as “secretory progenitors”, as they resemble intestinal label-retaining cells that are secretory progenitors. Two clusters (C6 and C30) were excluded from further analysis due to their high expression of hypoxic and apoptotic genes, leading us to classify them as hypoxic dying cells. Additionally, two other clusters (C34 and C35) contained too few cells (only 130 cells combined) and were excluded from further analysis, as their sample-specific nature might not accurately represent general cell behavior in gastric/IM cell development.

We constructed three-dimensional cell lineage trajectories using diffusion map analysis (Destiny 3.2, R package), which estimates diffusion-like dynamics within differentiating cells based on their distance metric (diffusion distance).^18^ ^19^ In short, this analysis utilized the normalized expression matrix to calculated the cell-to-cell transition probabilities, and produced eigenvectors of the diffusion components. The results were visualized by rgl (version 0.109, R package) and presented as interactive 3D graphics illustrating the transition relationship between cells. We pooled 4 AO/BO samples (GX023-BO, GX036-AO, GX059-BO, GX072-AO, Wnt+/-) and 5 IMO samples with high IM purity (GX001-IM3O, GX059-IMO, GX072-IMO, GX076-IMO, GX091-AgO^IMO^, Wnt+/-) to generate 3D lineage trajectories representative of gastric or IM cells cultured with or without Wnt3A (figure 2 and online supplementary figure 4D). We removed outlier cells that were distant along a diffusion component without a well-defined differentiation path, as they could represent stand-alone cell groups originating from rare subclones. Our results revealed two major branches in both gastric and IM cells, representing the gastric mucous/intestinal enterocyte trajectory and the secretory lineage trajectory, including enteroendocrine differentiation. We then analyzed each trajectory arm separately by constructing the stem-to-gastric mucous/enterocyte trajectory and the stem-to-secretory progenitor trajectory for the AO/BO or IMO pooled samples, respectively.

### Pseudotime inference of cell differentiation trajectories

To infer cell differentiation trajectories, we used scRNAseq data to perform pseudotime analysis and visualized the pseudotime values on 3D diffusion maps. First, we used Monocle 3^20^ to construct cell trajectories using Wnt+ and Wnt-sample pairs for each organoid, that captured the cell differentiation transition from growth medium to differentiation medium. Next, we used a stochastic RNA velocity model implemented in scvelo (v 0.2.4) to infer the direction of the trajectories.^21^ The velocity vectors were plotted and visualized as locally averaged vector fields on the trajectory-aligned cell populations. Latent time, a continuous variable, represents the relative progression of individual cells along their differentiation trajectory. It is an inferred measure that helps to identify the order of cellular events and transitions in a cell population, enabling the construction of a continuous trajectory of cellular differentiation. To estimate the latent time of cell trajectories in an unsupervised manner, we employed the CellRank (v1.5.1) method,^22^ which avoids artificial assignment of start and end points that may need prior knowledge of cell fates. This method uses a Markov chain approach to combine the velocity-based cell-to-cell direction vectors and the similarity-based diffusion map into directed cell lineages, revealing the latent time underlying cell trajectories. Finally, we integrated the latent time values from each individual Wnt+/-pair into pooled sample 3D diffusion maps to highlight the cellular differentiation trajectories.

### Calculation of fetal and YAP stemness signatures

Fetal and YAP stemness signature scores were calculated using the Single-Cell Signature Scorer tool in the Single-Cell Signature Explorer package.^13^ The genes comprising the fetal gene signature were derived from a study by Chen et al.,^23^ which curated a gene list based on previous studies^24^ ^25^ that identified a common set of genes that were up-regulated in the mouse fetal intestine (reminiscent of endoderm) compared with more advanced developmental stages. Chen et al. observed this fetal gene signature in colonic serrated lesions that showed concurrent loss of colon specific transcription factor and expression of gastric genes, inferring gastric metaplasia and reversion to a fetal state. This fetal signature was subsequently found in a YAP-dependent regenerative process in the mouse intestine under various forms of injury, including during helminth infection,^26^ inflammation^27^ or following irradiation.^28^ The genes comprising the YAP stemness signature were thus derived from the latter study that identified YAP-induced genes in a YAP mouse model showing Yap-dependent reprogramming of Lgr5(+) stem cells under radiation injury.^28^ Selection of the intestinal YAP stemness signature genes was accomplished by performing differential expression gene (DEG) analysis on the mouse data, comparing YAP doxycycline-induced overexpression in intestinal cells (DOX OE) or YAP knockout (KO) with their corresponding controls. Genes with YAP DOX logFC>1 and YAP KO log2FC<-1 were considered to be increased YAP stemness genes. Seven genes in the fetal and YAP stemness signatures showed overlap with the stomach signature and were removed from our analysis (online supplemental table 8).

### Cell fate tracking

We investigated gastric cell plasticity by analyzing cell fate diversity, which measures the potential for cells to differentiate into various cell lineages. We used the HPA single cell collection^29^ to analyze the normalized gene expression of 81 distinct cell types from normal tissue cells. We constructed a TPM gene expression matrix of all genes in the 81 cell types, and performed cell type specific gene selection. For each gene, we selected the 2 cell types with the highest expression levels to serve as representative cells for that gene. A gene was considered to be cell type-specific for a particular cell type when the gene expression TPMs in the top cells were greater than 10 and the log2 fold change against the 90-percentile expression in all the remaining cell types was larger than 3. We excluded the common genes shared between different cell types by comparing their expression against the average specific expression of the other cells. We then calculated the contribution coefficient for each gene on each cell type. Finally, the cell fate score for any single cells can by computed by the cell coefficient model, and then normalized into the cell fate probabilities, sum of which equal to 1. We estimated the cell plasticity by calculating the Shannon entropy of the 81 cell fate ratios into the cell diversity score, which were shown to be a good predictor of cellular pluripotency and plastic cell phenotype.^30^ The higher cell diversity means the more uniform distribution of different cell fates. The lower cell diversity means the cells are converged into a defined destiny. We examined the fetal and YAP stemness scores and cell fate diversity scores along the inferred pseudotime generated in the previous section, to track the plasticity dynamics during cell differentiation. Finally, we regressed the cell diversity into the Valley-Ridge (VR) space^31^ and projected our single cells on the 3D rendering diversity mountain by Generalized Additive Models (gam, version 1.22), where the DCs (diffusion components) and latent time were taken as the x-y bases and cell fate diversity entropy as the mountain height.

### Constructing normal and IM developmental trees using scRNAseq data for tracing cancer evolution in human IM and cancer tissues

We constructed development trees representing the differentiation trajectories of the gastric and IM organoid cells using an evolutionary clock model. Cells from AO/BO and IMO lineages were first grouped according to their velocity-inferred differentiation time (latent time). Cells within the same time bin (1/20 of the total time for each bin) and of the same cell type were clustered together as tree nodes. Subsequently, we calculated the molecular distance matrix between nodes based on the Spearman correlation of their batch-corrected expression profiles. Using the unweighted pair group method with arithmetic mean method (UPGMA), we constructed the development trees based on the node distance matrix and then fitted them to their latent time.^32^ After obtaining the tree-like architecture for the AO/BO and IMO lineages, we summarized the expression profiles in the terminal tree nodes (tips) using the node gene expression, while the internal node (ancestors) expression profiles were represented by the edge-weighted summation of branch lengths, with the terminal nodes contributing more to their ancestors’ profiles when they were closer. To explore the association between the evolutionary dynamics of our AO/BO or IMO organoids and gastric cancer evolution at the human tissue level, we referred to a recently published scRNA dataset comprised of gastric cancers and paired non-neoplastic gastric tissue.^33^ We focused our analysis on the epithelial cells after filtering out tumour microenvironmental cells, and only considered samples with an adequate number of epithelial cells for analysis (n>500). We performed cell clustering in each sample, examined their marker gene expression, and observed the presence of cell clusters exhibiting marker gene expression consistent with IM transition in some normal samples. The 7 normal samples derived from the gastric body or antrum were then sorted into IM cell clusters (average *CLDN3* expression >=0.5) and gastric cell clusters (average *CLDN3* expression <0.5). We correlated the expression profiles of each normal gastric or IM cell cluster with the nodes in our AO/BO and IMO lineage trees. The top most highly correlated interactions (Spearman correlation >0.75) between the tissue cell clusters and the organoid tree nodes were visualized using circus-tree plots. For gastric cancer tissues, we selected scRNAseq data from 5 tumours with high *CLDN3* expression (average *CLDN3* expression >0.5), indicating a potential origin from IM, for analysis. We generated cell clusters for each tumour and correlated their expression profiles with the nodes in our AO/BO and IMO lineage trees, displaying the top most highly correlated interactions (Spearman correlation >0.75) in the circus-tree plots. Finally, we calculated fetal and YAP stemness scores for each cell and compared them between normal gastric cells, IM cells and gastric cancer cells using the Mann-Whitney test.

### Differential gene expression analysis using scRNAseq data

Differential gene expression analysis was performed using the ‘FindMarkers’ function in Seurat (Version 3.2.3, R package)^15^ to investigate differential expression between IM cells in parental organoid and spheroid pairs, based on a non-parametric Wilcox rank sum test (after filtering out the gastric normal cells defined above from both groups). The up-and down-regulated genes were filtered according to log fold change > 0.4 or <-0.4 (fold change > 1.5 or < 0.666), respectively, and the adjusted *p*-value < 0.01, and cell proportion difference > 0.2. Then, differentially expressed genes observed in at least 2 out of 3 samples were chosen as common DEGs and ranked by the percentage of cells in spheroids showing up-regulation of gene expression. Gene function enrichment analysis was conducted separately on the up-/down-regulated genes in each IM spheroid and organoid pair using a hypergeometric test on the gene sets from the HALLMARK database,^34^ with the level of significance set at p<0.05.

### Identification of tumor-like cells using scRNAseq data

Single-Cell Identification of Subpopulations with Bulk Sample Phenotype Correlation (Scissor) method^35^ was applied to identify cancer phenotype-associated subpopulations using our scRNAseq data. This method integrates bulk RNAseq expression data with scRNAseq data by quantifying the similarity between each bulk sample and each single cell, followed by optimizing a regression model on the correlation matrix with the sample phenotype to identify relevant subpopulations. First, we constructed a gastric cancer signature based on the bulk RNAseq data from 38 normal organoids and 43 gastric tumor organoids from our previous study.^4^ Then, we used Scissor to apply a logistic regression model to optimize the expression correlation between each single-cell and bulk sample pair with the sample phenotype (tumor/normal). The sign of the regression coefficient indicates the relationship between the Scissor selected cells with the phenotype, such that cells with positive or negative coefficients are positively or negatively associated with a tumor-like phenotype, respectively.

### Single-cell analysis of copy number variations (CNVs)

InferCNV (R package, version 0.99.0, https://rdrr.io/bioc/infercnv/) was used to predict CNVs based on our scRNAseq data. A total of 1,500 single cells were randomly selected from GX059-BO and H019-NO as normal references for gastric and intestinal samples, respectively. Briefly, gene expression intensities at specific genomic positions (segments) were smoothed, and the average was calculated to represent the chromosomal copy numbers in comparison with the average of reference “normal” cells.

Furthermore, GX001-IM3O was subjected to single-cell copy number variation (scCNV) DNA sequencing. Single cells were prepared, as described, above using a cell sorter, encapsulated in cell beads, and lysed using the Cell Bead Polymer Kit (10xGenomics). The denatured genomic DNA was encapsulated, barcoded, and amplified for library construction using the Chromium Single Cell DNA Library and Gel Bead Kit (10xGenomics), according to the manufacturer’s instructions. The constructed single-cell DNA library was sequenced using the NovaSeq 6000 platform (Illumina). Raw sequencing reads were demultiplexed, aligned to the human genome (GRCh38), and scCNVs were called using Cell Ranger DNA (version1.0.0, 10xGenomics). The scCNV profile of GX001-IM3O was visualised using the Loupe scDNA Browser (version1.1.0, 10xGenomics).

### DNA methylation array and data analysis

DNA Methylation array was performed on 28 IM, 35 normal gastric and 4 normal intestinal organoids (online supplemental table 2). An average of 500ng of genomic DNA per sample was bisulfite converted using the EZ DNA Methylation kit (D5001-Ver.1.2.6, Zymo Research). The bisulfite-converted genomic DNA was then amplified, fragmented and precipitated. The re-suspended DNA was subsequently hybridized to probes attached to Infinium MethylationEPIC BeadChips (Infinium MethylationEPIC BeadChip Kit, version 1, Illumina), and the un-hybridized DNA was removed. Finally, the attached probes were subjected to single-base extension and stained with multiple antibodies. The beadchips were imaged and scanned using the Illumina iScan System. The data was analysed using GenomeStudio® (version 2011.1, Illumina) and GenomeStudio® Methylation Module (version 1.9, Illumina). Unsupervised hierarchical clustering was performed on the methylation array data from 67 organoids after BMIQ normalization and filtering away CpG probes with N/A values across the cohort. Differential methylation analysis was performed to identify methylation differences between a “Consensus IM” and “Consensus Normal” group using an empirical Bayes moderation method^36^ implemented in limma (R package, version ?). The “Consensus IM” group consisted of 12 IM organoids in the IMOme cluster 1 (online supplemental figure 7A) that overlapped with the IMOexp cluster 1 (figure 1A). The “Consensus Normal” group consisted of 31 normal organoids that were classified as normal by gene expression. CpG probes with a median difference in beta-values >0.3 or <-0.3, and adjusted *p*-value<0.01 were considered to be hypermethylated or hypomethylated in IM, respectively. Finally, methylation regulating events were identified by analysing the correlation between the methylation profile of promoter CpGs and their corresponding gene expression (Pearson correlation coefficient < -0.6, *p*-value < 0.01).

To measure the global deviation in DNA methylation patterns from a normal reference, we calculated the Hyper-Z index, which measures aberrant hypermethylation over promoter CpG islands in a given sample, and the Hypo-Z index, which measures aberrant hypomethylation over opensea probes (intergenic regions of low CpG density), for each sample.^37^ Briefly, for each sample, an index was calculated by averaging the Z scores (relative to the mean and standard deviation of normal samples) across all probe clusters in a genomic region: promoter CGIs for the Hyper-Z index and opensea probes for the Hypo-Z index. Since promoter CGI regional clusters are usually unmethylated in normal tissue, only regions for which the Z score is positive contribute to the Hyper-Z index. In contrast, for open sea regional clusters, which are usually methylated in the normal tissue, only regions for which the Z score is negative contribute to the Hypo-Z index. Thus, for each sample, we calculated an overall Hyper-Z index by summarizing all the positive Z-scores and a Hypo-Z index by summarizing all the negative Z-scores, reflecting the global level of aberrant DNA hypermethylation and hypomethylation in that sample, respectively.

### Whole exome sequencing and data analysis

WES was performed on 70 gastrointestinal organoids (28 IM and 42 normal stomach), 5 IMOLs, and 3 IM spheroids (online supplemental table 2). A total of 550ng of genomic DNA per sample, after fragmentation using the Covaris S2, was input for library preparation using the KAPA Hyper Prep Kit (KR0961-V1.14, Biosystems), following the manufacturer’s protocol. 330ng of library DNA each, from 12 samples, were pooled for exome capture using xGen Lockdown Probes and Reagents (IDT), according to standard protocols. The captured libraries were sequenced using either 151bp paired end reads on the NovaSeq6000/HiSeqX Ten or 101bp paired end reads on the HiSeq1500 (Illumia). The mean coverage for the organoids and germline DNA were 50X. The sequencing reads were aligned against the human genome (hg19) using BWA (version 0.7.10)^38^ and samtools (version 0.1.19).^39^ The duplicated reads in the BAM files were removed using Picard (version 1.73) (http://broadinstitute.github.io/picard/) and recalibrated by Genome analysis toolkits (GATK) (version 3.2-2) with default settings.^40–42^ Similar to our previous studies,^4^ ^5^ SNVs and INDELs were called by the GATK UnifiedGenotyper function based on the following criteria,

1. stand_emit_conf of 10

2. stand_call_conf of 50

The remaining variants were filtered using the GATK VariantFiltration function based on the following criteria,

1. Quality of depth of 2

2. Fisher strand score 60 (SNVs) and 200 (INDELs)

Somatic mutations for each sample were identified by comparison with their paired germline DNA from blood leukocytes, with SNVs detected by Strelka^43^ and indels detected by the GATK SomaticIndelDetector function. The list of somatic mutations was finalised by comparing results from the two different software packages.

DNA copy number analysis was performed by cnv_facets (version 0.14.0),^44^ with default settings as previously described.^4^ The organoids with chromosomal aberrations were identified based on the following parameters,

1. DNA segment size > 10 million bp

2. Estimated cell fraction per sample > 10%

3. Percentage of chromosome length with alterations > 30%

### Cell proliferation assay

IM and normal gastric organoids were trypsinized into single cells and 10,000 cells in 50μL Matrigel were seeded into a 24-well plate, in triplicate, as previously described.^4^ Cells were cultured in a standard human gastric organoid culture medium containing 10µM Y-27632 for 7 days. Subsequently, the organoids were dissociated into single cells again and the total number of live cells was counted by trypan blue exclusion using a LUNA™ Automated Cell Counter (Logosbio). The cell growth rate was calculated as an average of 3 replicates using the following equation:

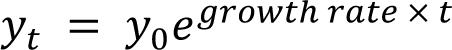

(*yt* = number of cells at final time point, *y0* = number of cells at initial time point, t = time in hours). The mean doubling time (hours) was calculated as

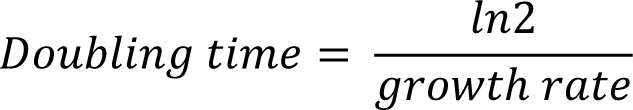

### Cell viability assay

IM organoids were dissociated into single cells and 5,000 cells in 50µL Matrigel were plated into a 24-well plate, in triplicate. The cell viability of organoids was accessed using the Cell Titre-Glo 2.0 Kit (Promega) upon FGF10 or ROCK inhibitor (Y-27632) withdrawal at day 6 or 15, respectively. The raw luminescent signals were obtained from a SpectraMax L microplate reader (Molecular Device). The cell viability percentage was calculated by normalization to gastric organoids grown in full human organoid culture medium.

### Growth factor withdrawal assay

Growth factor withdrawal assay was performed as previously described.^1^ An equal number of cells (n=5,000) for IM, normal gastric and normal intestinal organoids were plated onto each well of a 48-well plate in duplicate. Essential supplements, including Wnt-3A, Rspo1, Noggin, EGF, TGFβi (A83-01) and FGF10 were individually removed from the culture medium. The organoids were passaged at a 1:3 ratio every 7-10 days. The effect of withdrawing each growth factor or inhibitor on organoid growth was studied by comparing cell proliferation rates with controls grown in full human organoid medium on a weekly basis.

### Spheroid formation assay

To investigate cell-matrix dependence, IM and normal gastric organoids were first trypsinized into single cells. Twenty thousand cells were plated onto a 24-well ultra-low binding plate (Corning) in 5 replicates and cultured in suspension. Fresh stomach medium containing 10µM Y-27632 was added every 3-4 days. On day 14, the spheroids were centrifuged at 1,000 x g for 5 minutes and dissociated into single cells again. The total number of live cells was determined by trypan blue exclusion, counting cells from 3 replicates. In parallel, the remaining cells were continuously grown in suspension for anchorage independent cell selection.

### qRT-PCR and immunoblotting

Gene expression in our organoid cohort was studied by qRT-PCR using the QuantiFast SYBR Green PCR kit (Qiagen) and the LightCycler 480 Instrument II (Roche). All reactions were performed in triplicate, utilizing *GAPDH* expression as a normalization control. The primers used in this study are listed in online supplemental table 15.

For total protein extraction, organoids at 80-90% confluence were harvested using a lysis buffer (150 mM NaCl, 50 mM Tris pH 7.4, 1% NP40 & 1mM EDTA) supplemented with protease and phosphatase inhibitors (Roche), according to the manufacturer’s instructions. Protein lysates (60µg) were resolved on SDS-PAGE gels (10% acrylamide) and transferred to nitrocellulose membranes (Pall). The membranes were incubated with anti-CHGA (1:250, Sigma) or anti-GAPDH (1:2,000, Santa-Cruz) antibodies at 4°C overnight, and protein expression was detected using a western blotting detection reagent (Cytiva Amersham^TM^ ECL^TM^ Prime). The signals were captured by exposing the membranes to X-ray films, which were subsequently developed and scanned.

### Validation of CNVs by droplet digital PCR (ddPCR)

CNVs identified in gastric organoids by WES analysis were validated using the PrimePCR^TM^ ddPCR^TM^ Copy Number Variation Assay (BioRad). Briefly, 500ng of genomic DNA was digested in 10U of HaeIII (Thermo Fisher Scientific), at 37°C, for one hour. In parallel, DNA from the IM spheroids, which were cultured in suspension for over 4 weeks, was extracted using proteinase K. The spheroids were lysed in 10-15µL proteinase K (PK) buffer at 56°C overnight and subsequently heated at 95°C for 15 minutes. The resulting whole lysate was digested by HaeIII restriction enzyme as described above.

Next, 50ng of the digested genomic DNA or 2µL of digested genomic DNA extracted from PK buffer was added to a water-based reagent mixture and mixed thoroughly. Droplets were encapsulated using the QX200 Droplet Generator (BioRad), following the manufacturer’s instructions. Approximately 16,000-20,000 droplets were partitioned into a volume of 40µL per sample, transferred to a 96-well PCR plate and amplified through 40 PCR cycles using specific FAM and HEX-tagged probes (online supplemental table 15). The fluorescence intensity from each DNA-to-oil droplet was detected by a duplex-colour detection system on the QX200 Droplet Reader (BioRad). The dual signals’ intensity of each droplet was quantified using the QuantaSoft software (BioRad) and each droplet was classified into one of four clusters: reference^-^/target^-^, reference^+^/target^-^, reference^-^/target^+^ and reference^+^/target^+^. The average number of DNA copies of a sample was estimated using Poisson distribution modelling and the overall copy number was calculated with the following formula (assuming a normal reference chromosome has two DNA copies):

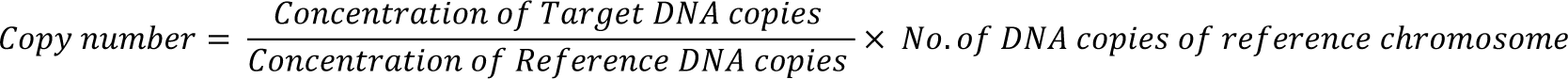

---

**Supplemental figure 1.**
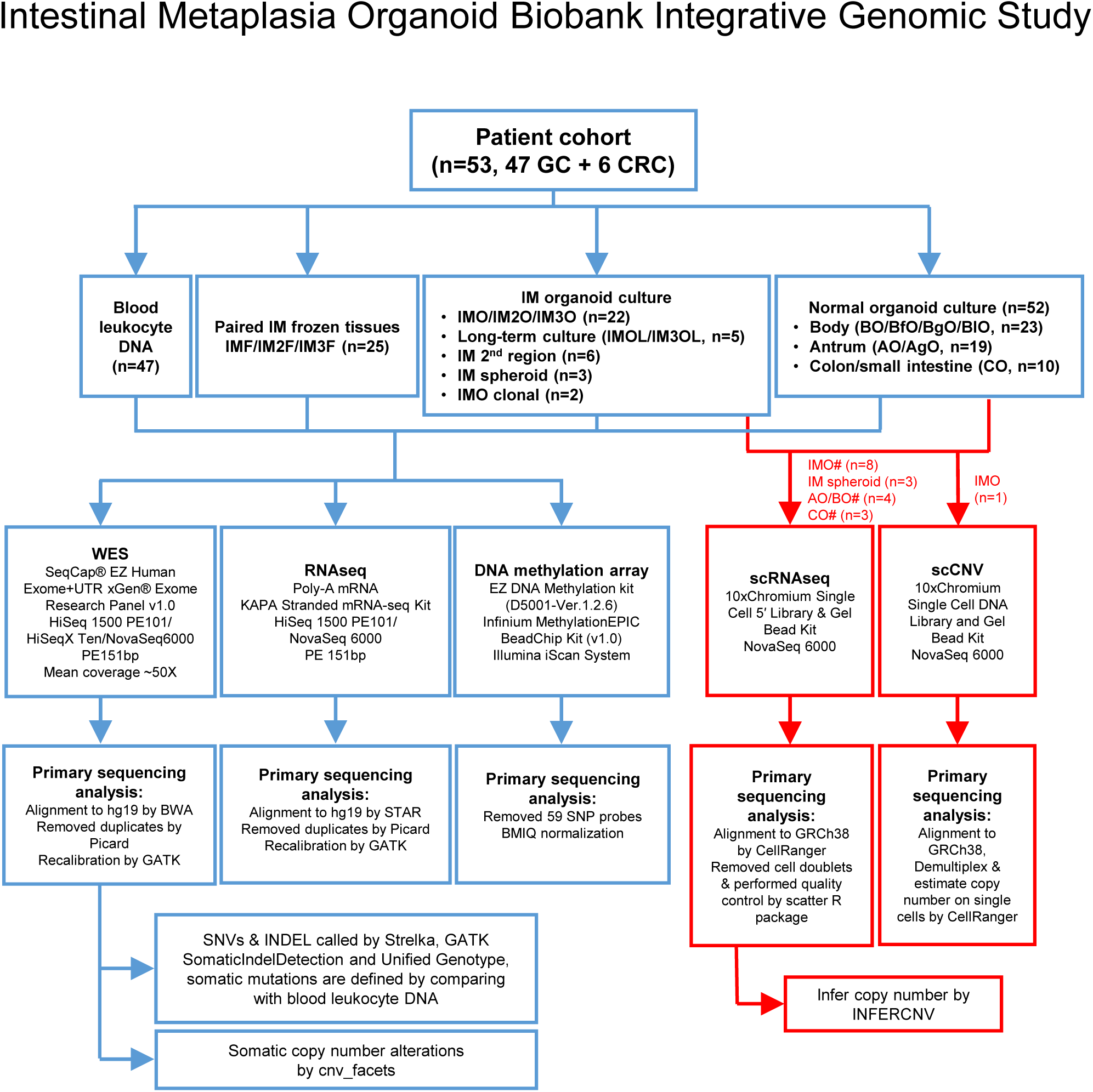
An overview of the IM organoid bank integrative genomic study. A flow chart summarizing the number of samples and analysis pipeline for the IM organoid biobank integrative genomic study. IM1F/IM2F/IM3F, IM frozen tissues; IMO/IM2O/IM3O, IM organoids; IMOL/IM3OL, IM organoids from long-term culture; BO/BfO/BgO/BIO, body organoids; AO/AgO, antrum organoids; CO, normal intestinal organoids; #, with paired differentiated samples. Blood leukocyte DNA was used as a reference for WES.

**Supplemental figure 2.**
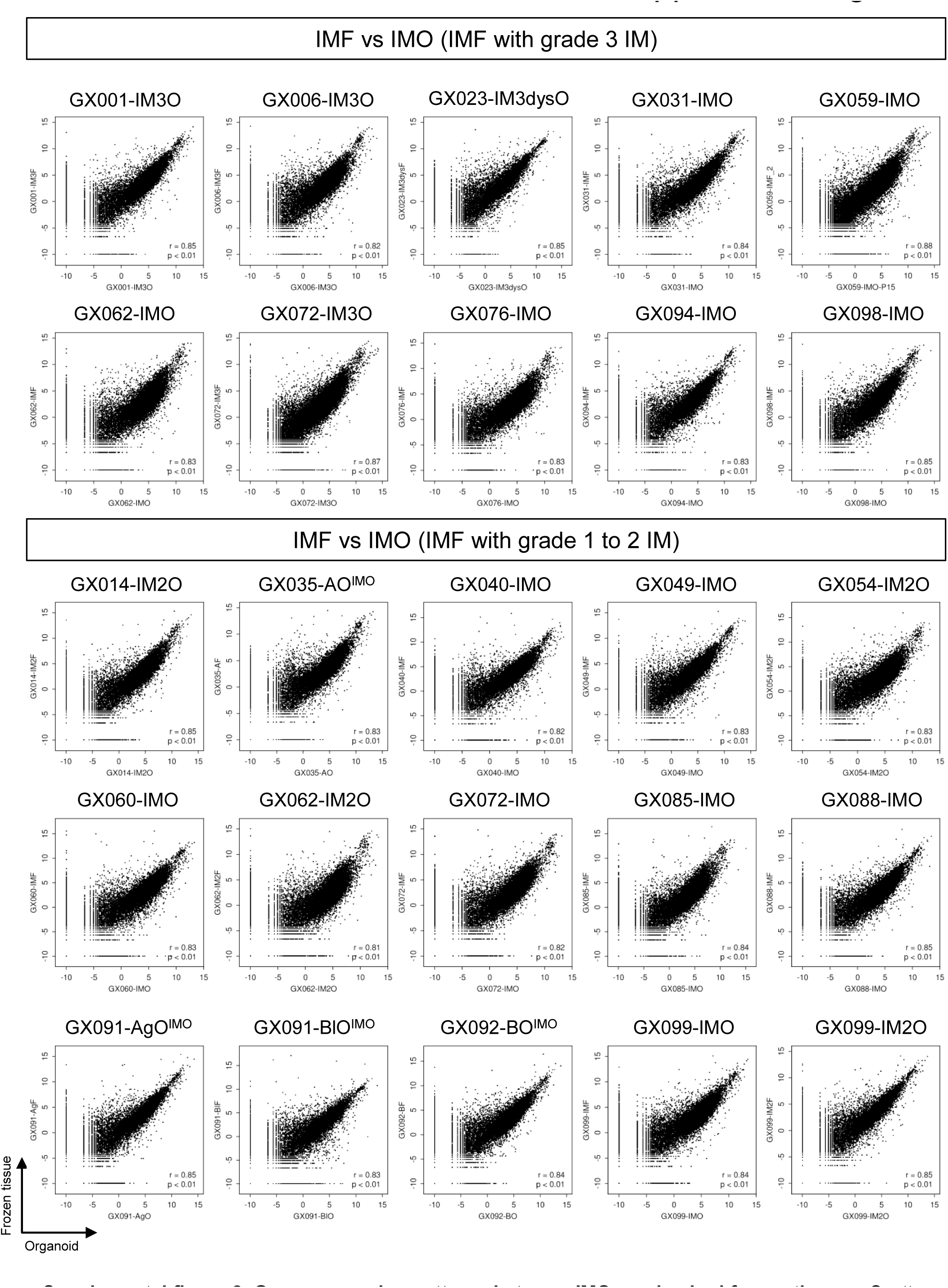
Gene expression patterns between IMOs and paired frozen tissues. Scatter plots illustrating that IM organoids (x-axis) recapitulated the gene expression profiles of the paired frozen tissues (y-axis) in terms of the log2 TPM values of individual genes. IMF/IM2F/IM3F, IM frozen tissues; IMO/IM2O/IM3O/IM3dysO, IM organoids; AgF/AF/BF, antrum or body frozen tissues.

**Supplemental figure 3.**
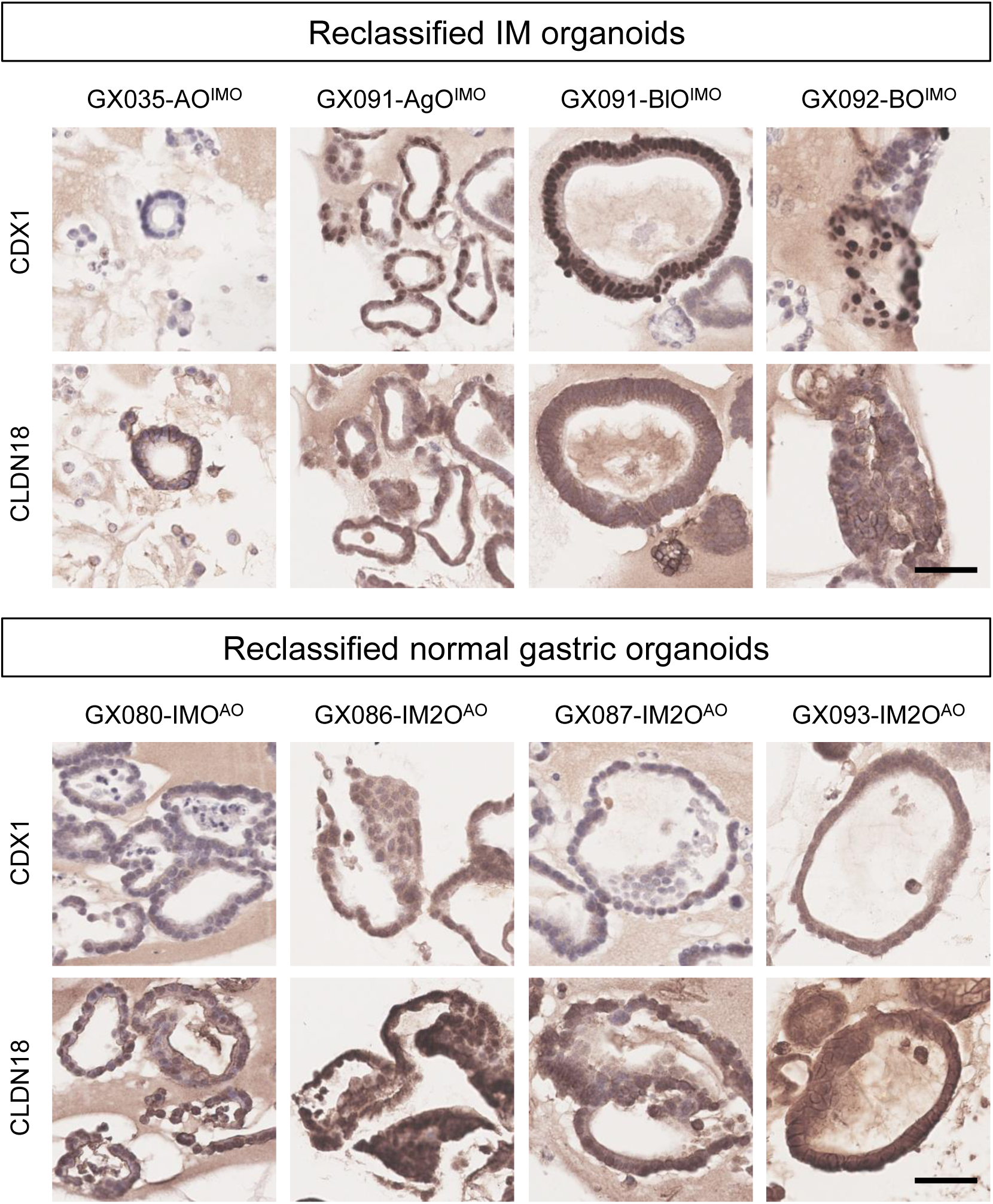
Protein expression of IM and gastric markers. IHC staining of CDX1 and CLDN18 in the reclassified IM (upper panel) and normal gastric (lower panel) organoids, related to Figure 1. Scale bar: 50µm.

**Supplemental figure 4.**
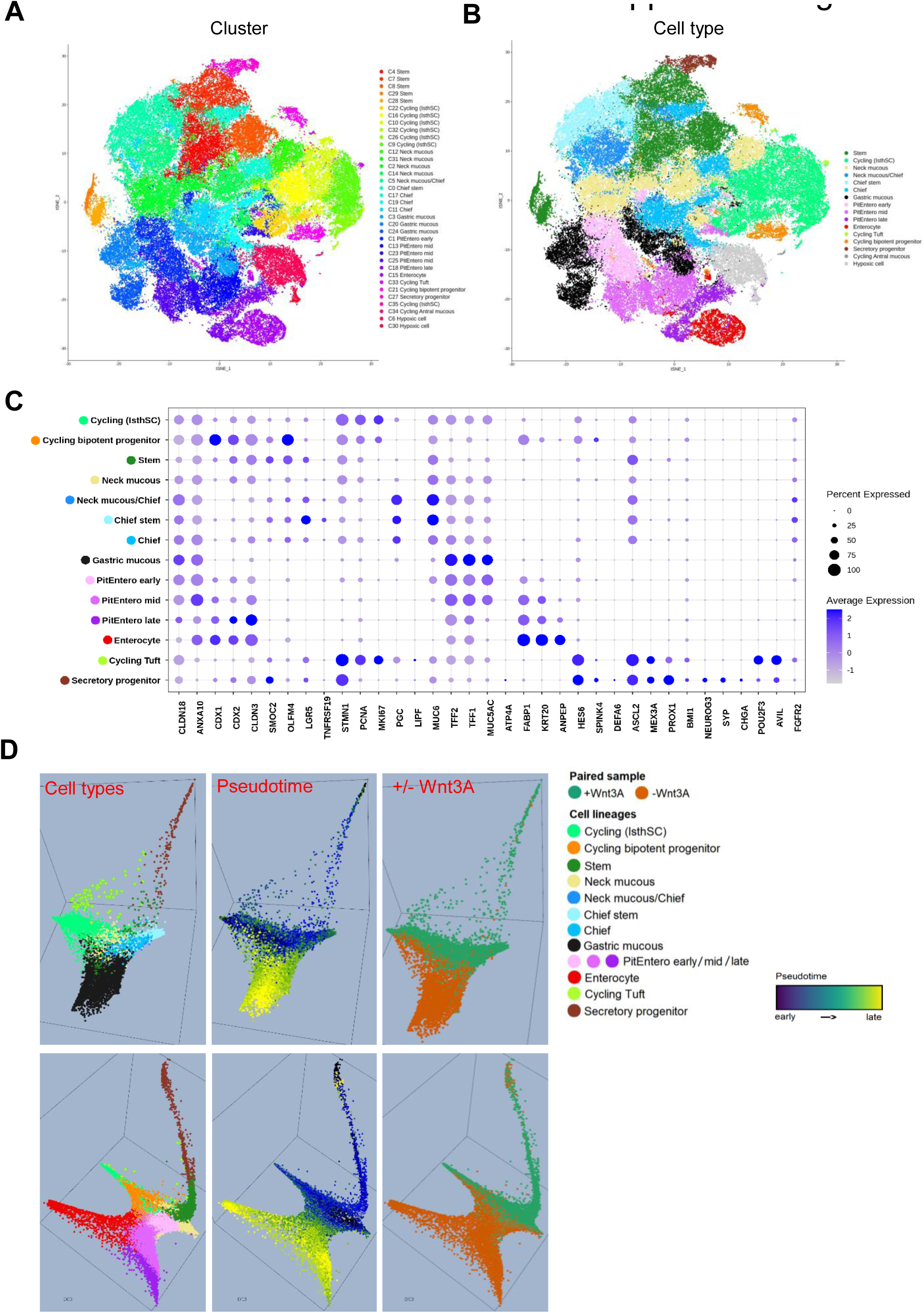
Single-cell RNA analysis of normal and IM organoids. (A-B) t-SNE plots showing an overview of the single cells after data integration. The cells are color-coded by cell clusters (A) and cell types (B). (C) Dot plot showing gene expression of specific stomach and colon lineage markers in 14 merged cell types. The dot size represents the fraction of cells that express each gene, and the color denotes the average expression level. (D) Diffusion map of AO/BO (top) and IMO (bottom) cell lineage trajectories.

**Supplemental figure 5.**
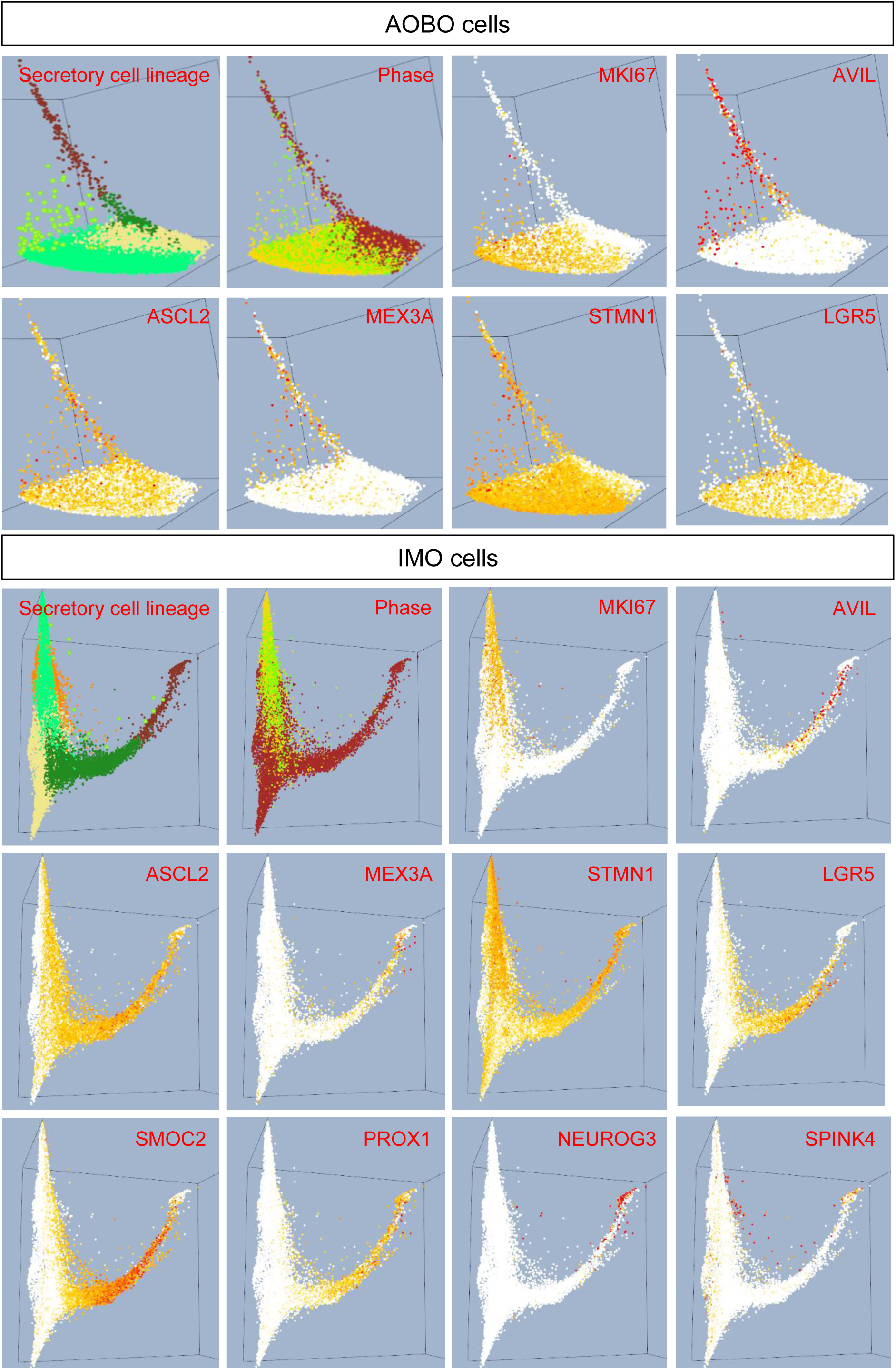
3-D diffusion map of secretory cell lineage from AO/BO sample integration and IMO sample integration, +/-Wnt-3A. Cells are colored according to cell lineage, as denoted in figure 3, or by the expression intensity of a specific gene, as labeled in each box.

**Supplemental figure 6.**
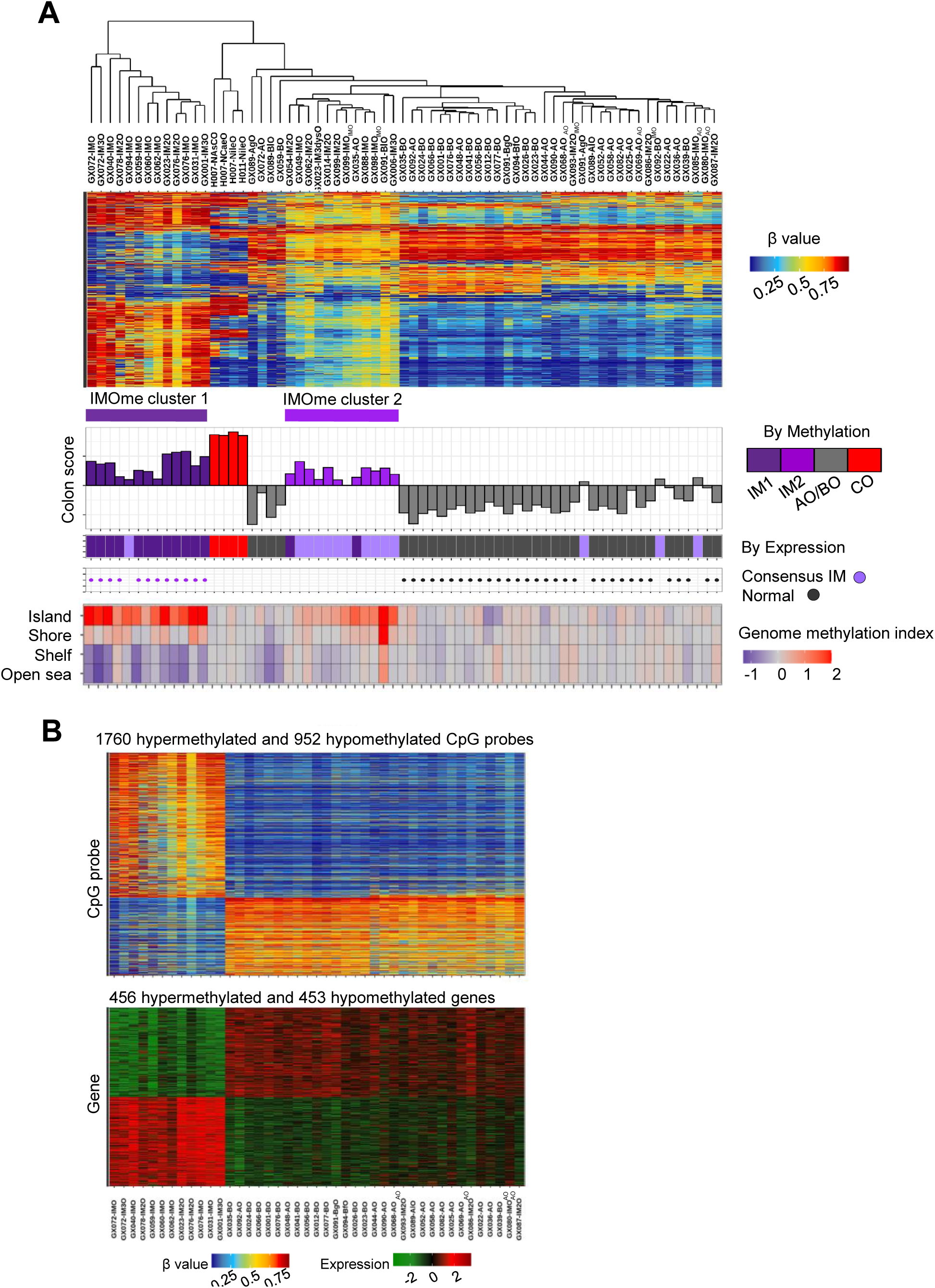
The epigenomic landscape of IMOs. (A) Hierarchical clustering using CpG probes segregated IMOs into 2 IMO methylation clusters that showed significant overlap with IMO expression clusters. Heatmap showing the differential methylation profiles at islands, shore, shelf and open seas of the organoids. Color bars represent the colon signature score for each sample from the bulk transcriptome analysis. (B) DMR analysis between consensus IM and normal organoids identified a list of negatively correlated CpG-gene pairs with their corresponding β-values (top) and normalized gene expression (bottom) presented as a heatmap. BO/BfO/BgO/BlO, body organoids; AO/AgO, antrum organoids; CO, intestinal organoids; IMO/IM2O/IM3O/IM3dysO, IM organoids.

**Supplemental figure 7.**
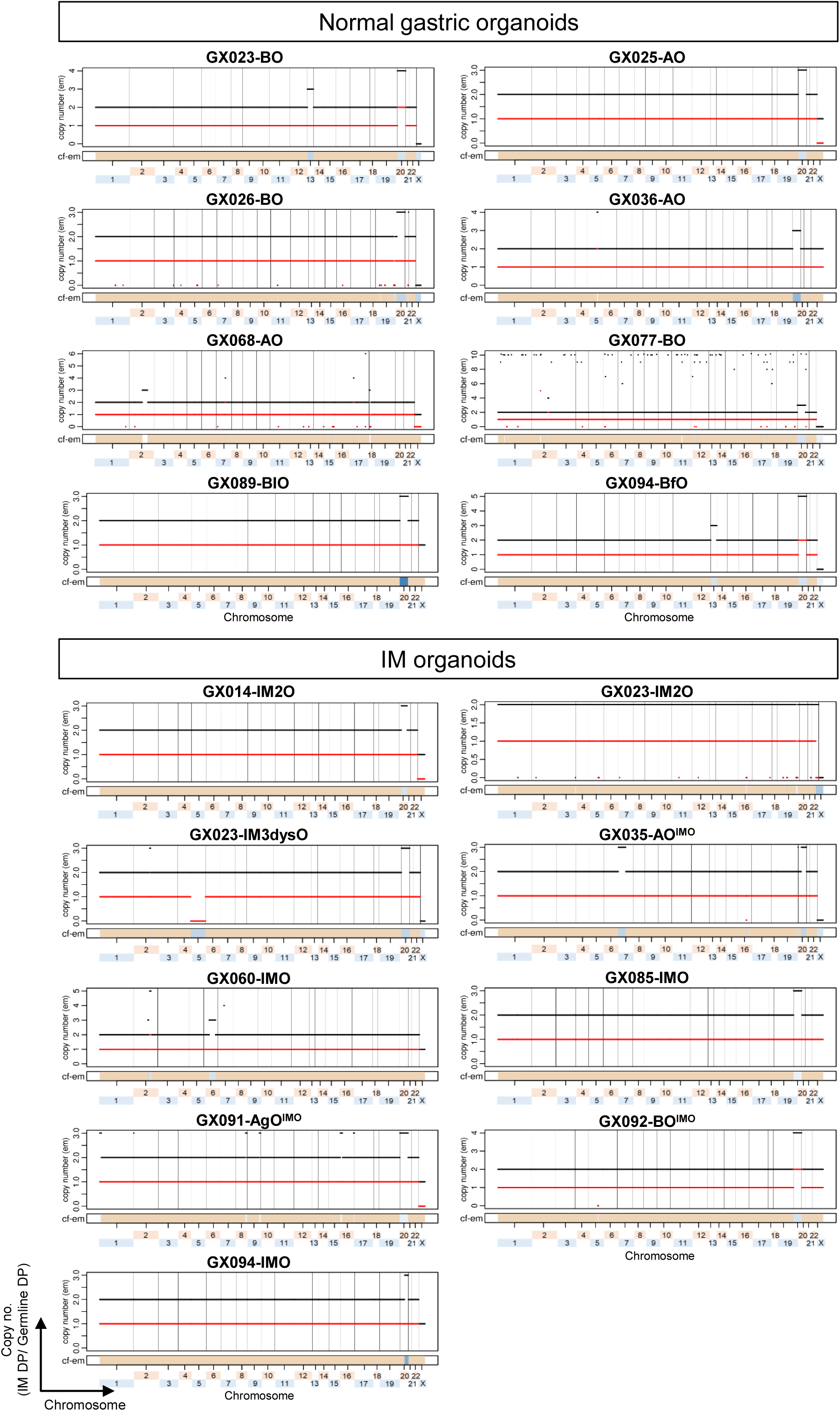
Copy number plots showing the CNVs detected in normal gastric and IM organoids. Black and red lines indicate the total and minor copy numbers, respectively.

**Supplemental figure 8.**
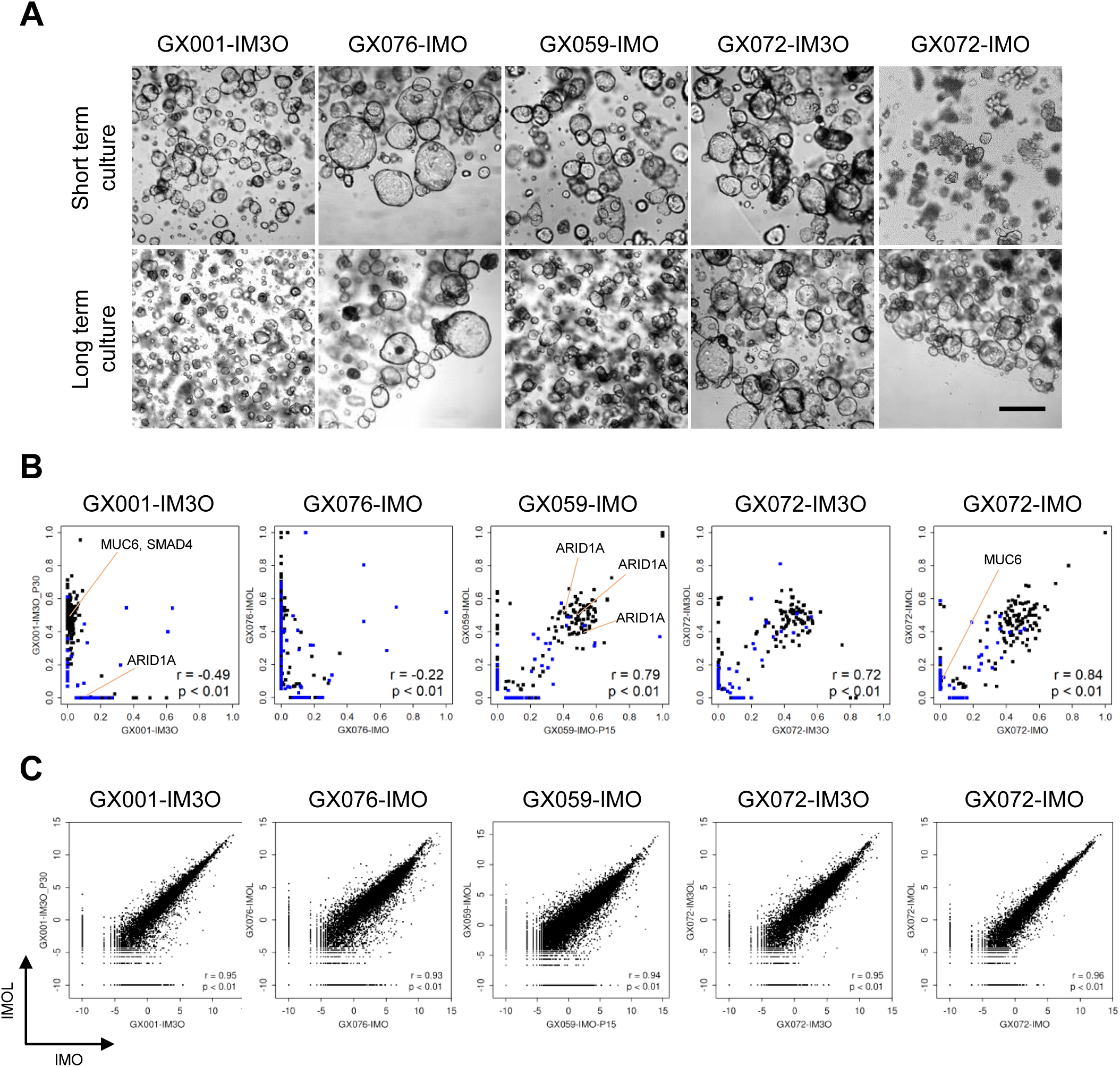
The morphology, genomic and transcriptomic stability of IMOs between short- and long-term cultures. (A) Bright field images showing the indistinguishable morphology of 5 IMOs before and after long-term culture. Scale bar: 500µm. (B) The presence of clonal dominance in GX001-IM3O, GX059-IMO and GX076-IMO, as well as the clonal features of GX072-IM3O and GX072-IMO were stably maintained *in vitro* after long-term culture. Scatter plots illustrating the mutant allelic fractions of SNVs in IMOs between short- (x-axis) and long-term (y-axis) cultures. (C) Scatter plots illustrating the concordant transcriptomic profiles of 5 IMOs between short- (x-axis) and long-term (y-axis) cultures by RNAseq. IMO/IM3O, IM organoids; IMOL/IM3OL, long-term cultured IM organoids.

**Supplemental figure 9.**
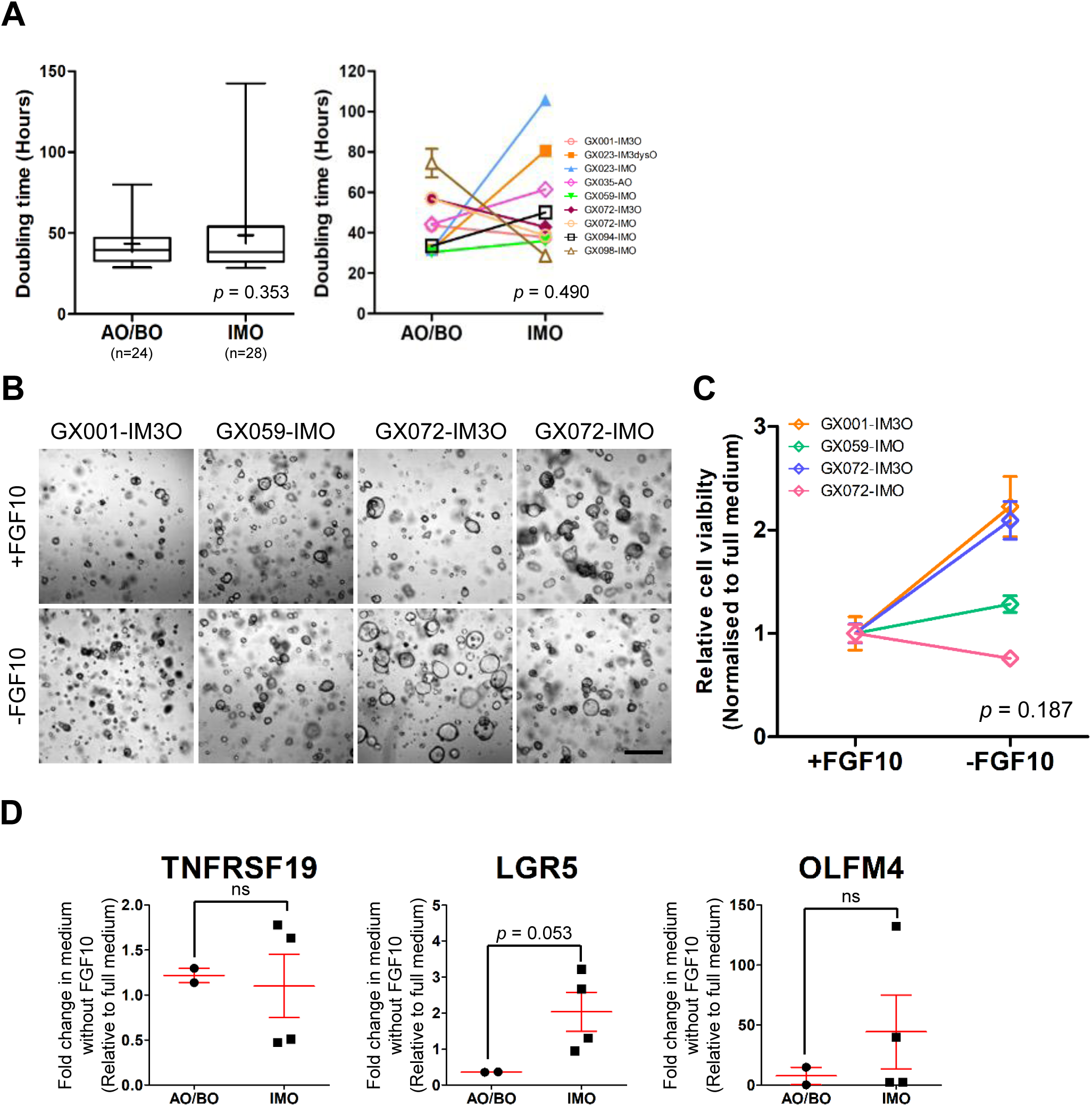
The growth rate of organoids, and characterization of IMOs in FGF10-free culture medium over 6 months. (A) The growth patterns of AOs/BOs and IMOs in terms of the doubling time (left). The whiskers indicate the slowest and fastest doubling times, the horizontal lines show the median, and the “+” indicates the median and mean values, respectively (left). P value obtained by student’s t-test. The doubling times of 9 IMO and AO/BO pairs were compared (right). Error bars indicate mean ± SD from three technical replicates. Paired student’s t-test was performed between two groups. (B) Brightfield images of 4 IMOs cultured in human stomach medium, with or without FGF10 after 6-months in culture. Scale bar: 500µM. (C) The relative cell viability of 4 IMOs cultured under standard or FGF10 free conditions. Error bars indicate mean ± SD from three technical replicates. P value was calculated using a paired student’s t-test. (D) Dot plots illustrating the relative fold change by qRT-PCR of stem cell markers in IM and normal gastric organoids from an FGF10-free culture medium. Error bars indicate mean ± SD. ns, not significant; \**p* < 0.05; \*\**p* < 0.01; \*\*\**p* < 0.001.

**Supplemental figure 10.**
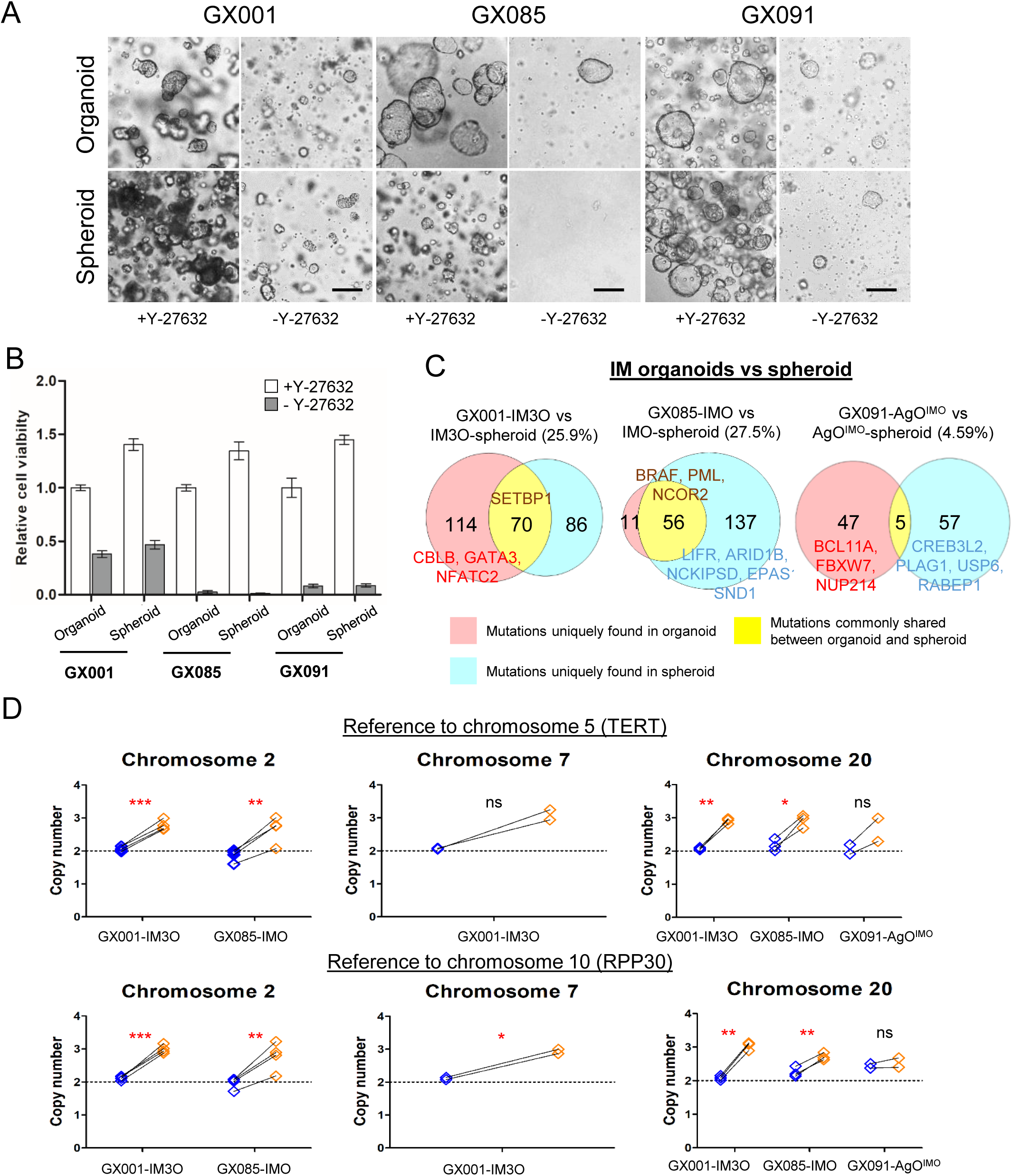
The reliance on Y-27632, genomic differences, and detection of copy number changes by droplet digital PCR (ddPCR) between IM organoid and spheroid pairs. (A) Three pairs of IM organoids and spheroids were cultured in Matrigel and underwent selection. All three were Y-27632 dependent, with significant cell death observed. Bright field images were taken at Day 15. Scale bar: 100µm. (B) Bar chart showing the cell viability of 3 IMOs and spheroids upon Y-27632 withdrawal (denoted as –Y-27632) at Day 15, relative to the corresponding IMOs cultured in full human stomach medium (denote as +Y-27632) before spheroid selection. (C) Venn diagrams showing a low percentage of shared somatic SNVs between organoids and spheroids. Only the top 5 COSMIC genes with protein altering SNVs in terms of allelic fraction in the IM spheroids or corresponding organoids were labelled if there were more than 5 SNVs. (D) The copy number measured by ddPCR in organoids (blue) and corresponding spheroids (orange) after culturing in the ultra-low binding culture plates for 4-6 weeks. The genes, *TERT* and *RPP30*, located on chromosomes 5 and 10, respectively, were used as reference for data normalization. Statistics were performed by paired student’s t-test. ns, not significance; \**p* < 0.05; \*\**p* < 0.01; \*\*\**p* < 0.001.

**Supplemental figure 11.**
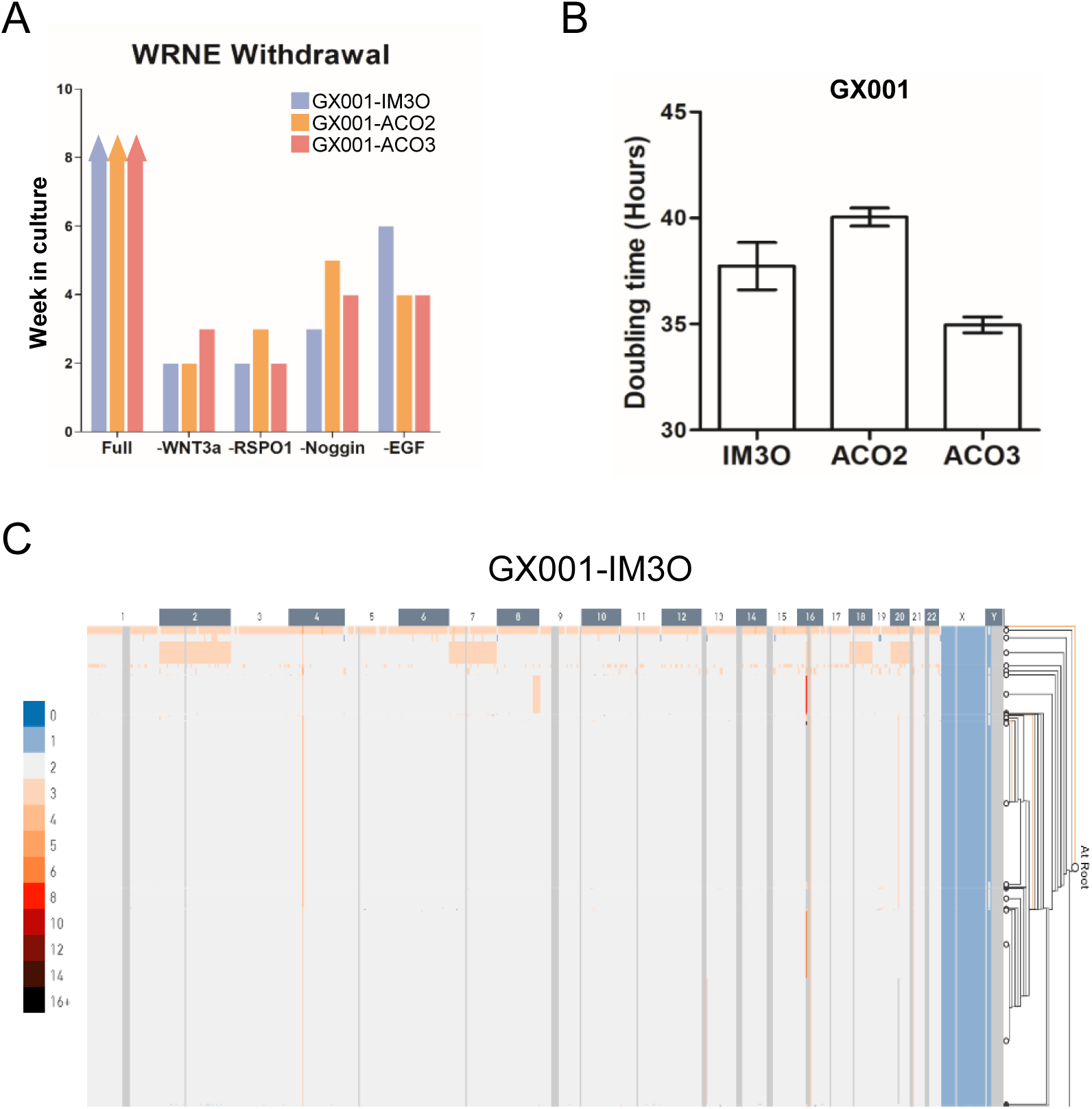
Characterization of clonal IM organoids and single-cell CNV profiling. A) Bar chart showing the weeks in culture of the IMOs upon individual growth factor withdrawal. Bars with an arrowhead denote organoids that could be cultured long-term (>6 months). (B) The growth behavior of pool and its clonal organoids, in terms of the doubling time. (C) Hierarchical clustering of single cells (rows) from GX001-IM3O based on the copy number quantitated by scCNV. A small subset of cells (n=64, orange colored) showed copy number gain of chromosomes 2, 7, 18 and 20 whilst most of the single cells were copy number neutral.

**Supplemental figure 12.**
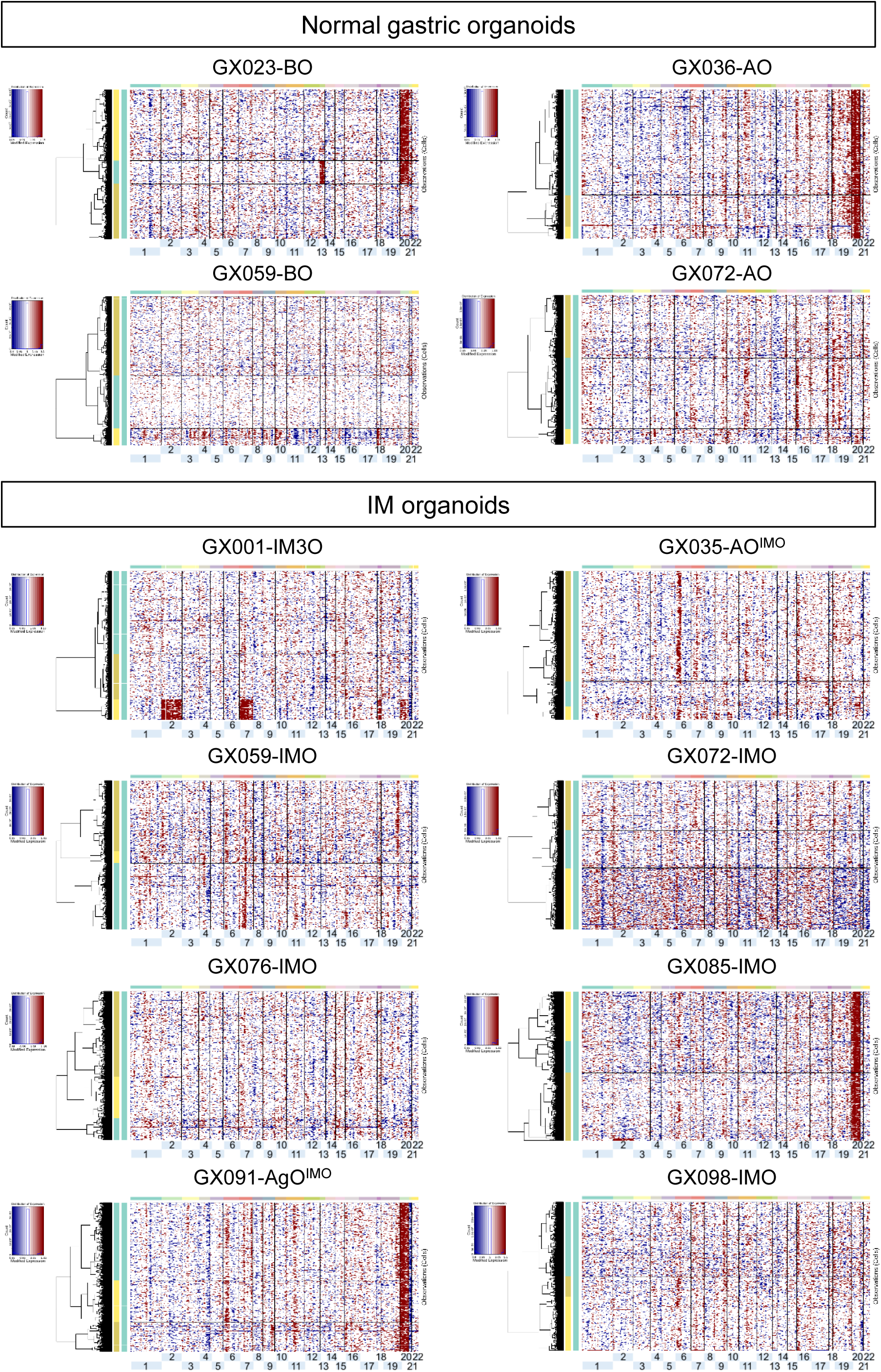
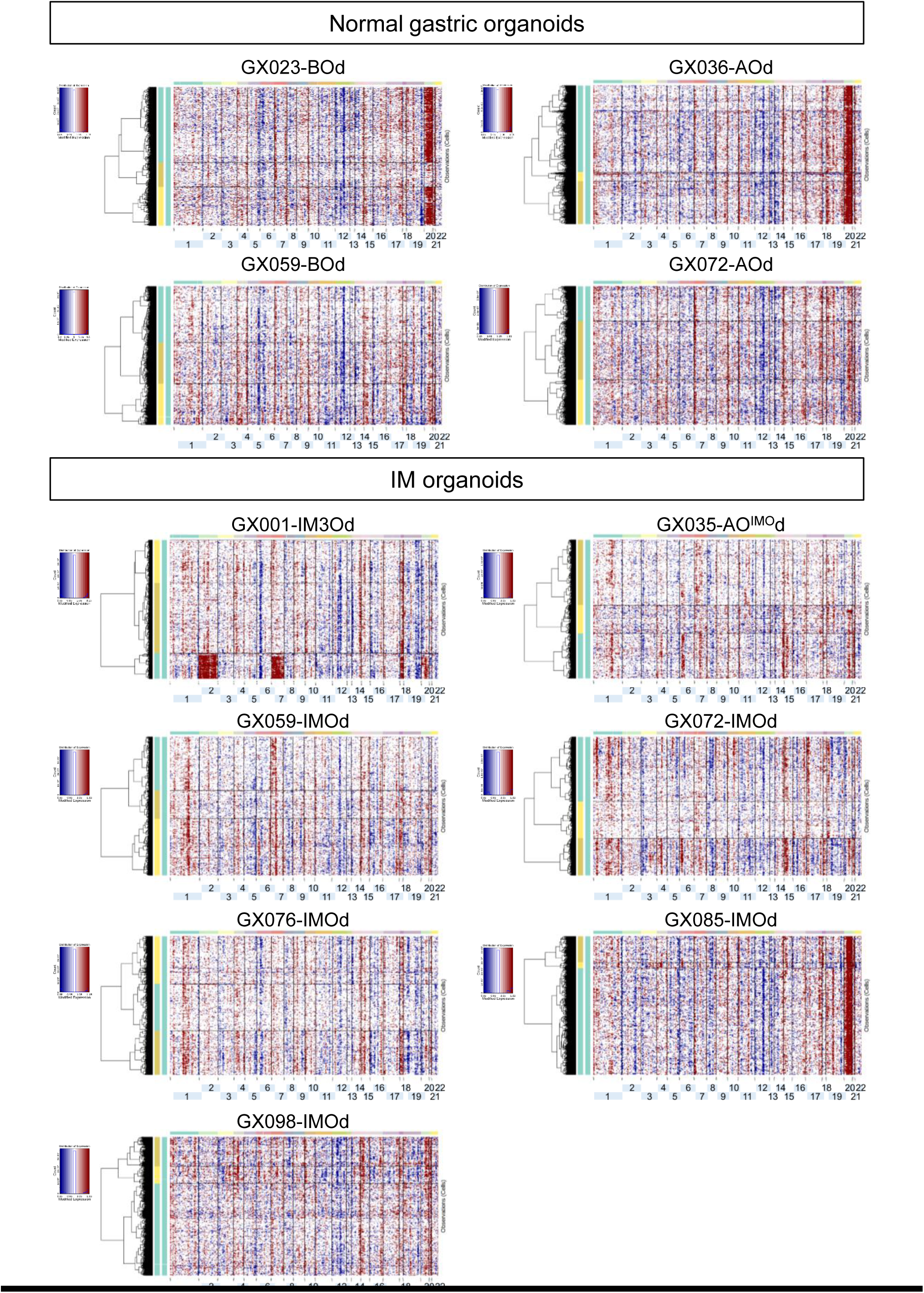
Hierarchical clustering of single cells (rows) from normal gastric and IM organoids based on the inferred copy number from scRNAseq data. The cells with copy number gain and loss are indicated as red and blue, respectively.

## REFERENCES

1 Mutoh H, Hakamata Y, Sato K, et al. Conversion of gastric mucosa to intestinal metaplasia in Cdx2-expressing transgenic mice. Biochemical and Biophysical Research Communications 2002;294:470–9.

2 Silberg DG, Sullivan J, Kang E, et al. Cdx2 ectopic expression induces gastric intestinal metaplasia in transgenic mice. Gastroenterology 2002;122:689–96.

3 Mutoh H, Sakurai S, Satoh K, et al. Cdx1 induced intestinal metaplasia in the transgenic mouse stomach: comparative study with Cdx2 transgenic mice. Gut 2004;53:1416–23.

4 Altayar O, Davitkov P, Shah SC, et al. AGA Technical Review on Gastric Intestinal Metaplasia-Epidemiology and Risk Factors. Gastroenterology 2020;158:732–44.

5 Gawron AJ, Shah SC, Altayar O, et al. AGA Technical Review on Gastric Intestinal Metaplasia-Natural History and Clinical Outcomes. Gastroenterology 2020;158:705–31.

6 Shiao YH, Rugge M, Correa P, et al. p53 alteration in gastric precancerous lesions. The American Journal of Pathology 1994;144:511–7.

7 Busuttil RA, Zapparoli GV, Haupt S, et al. Role of p53 in the progression of gastric cancer. Oncotarget 2014;5:12016–26.

8 Guo Y, Zhou J, Huang A, et al. Spatially defined microsatellite analysis reveals extensive genetic mosaicism and clonal complexity in intestinal metaplastic glands. Int J Cancer 2015;136:2973–9.

9 Lee J-H, Park S-J, Abraham SC, et al. Frequent CpG island methylation in precursor lesions and early gastric adenocarcinomas. Oncogene 2004;23:4646–54.

10 Huang KK, Ramnarayanan K, Zhu F, et al. Genomic and Epigenomic Profiling of High-Risk Intestinal Metaplasia Reveals Molecular Determinants of Progression to Gastric Cancer. Cancer Cell 2018;33:137–50.

11 Huang KK, Ma H, Chong RHH, et al. Spatiotemporal genomic profiling of intestinal metaplasia reveals clonal dynamics of gastric cancer progression. Cancer Cell 2023;41:2019–37.e8.

12 Kumagai K, Shimizu T, Takai A, et al. Expansion of Gastric Intestinal Metaplasia with Copy Number Aberrations Contributes to Field Cancerization. Cancer Res 2022;82:1712–23.

13 Takeuchi C, Yamashita S, Liu YY, et al. Precancerous nature of intestinal metaplasia with increased chance of conversion and accelerated DNA methylation. Gut 2023.

14 Wang K, Yuen ST, Xu J, et al. Whole-genome sequencing and comprehensive molecular profiling identify new driver mutations in gastric cancer. Nature genetics 2014;46:573–82.

15 Chen X, Leung SY, Yuen ST, et al. Variation in gene expression patterns in human gastric cancers. Mol Biol Cell 2003;14:3208–15.

16 Lee SH, Contreras Panta EW, Gibbs D, et al. Apposition of Fibroblasts With Metaplastic Gastric Cells Promotes Dysplastic Transition. Gastroenterology 2023;165:374–90.

17 Nowicki-Osuch K, Zhuang L, Cheung TS, et al. Single-Cell RNA Sequencing Unifies Developmental Programs of Esophageal and Gastric Intestinal Metaplasia. Cancer Discov 2023;13:1346–63.

18 Tsubosaka A, Komura D, Kakiuchi M, et al. Stomach encyclopedia: Combined single-cell and spatial transcriptomics reveal cell diversity and homeostatic regulation of human stomach. Cell Rep 2023;42:113236.

19 Zhang P, Yang M, Zhang Y, et al. Dissecting the Single-Cell Transcriptome Network Underlying Gastric Premalignant Lesions and Early Gastric Cancer. Cell Rep 2019;27:1934–47 e5.

20 Yan HHN, Siu HC, Law S, et al. A Comprehensive Human Gastric Cancer Organoid Biobank Captures Tumor Subtype Heterogeneity and Enables Therapeutic Screening. Cell Stem Cell 2018;23:882–97.

21 Yan HHN, Siu HC, Ho SL, et al. Organoid cultures of early-onset colorectal cancers reveal distinct and rare genetic profiles. Gut 2020;69:2165–79.

22 Lange M, Bergen V, Klein M, et al. CellRank for directed single-cell fate mapping. Nat Methods 2022;19:159–70.

23 Han S, Fink J, Jorg DJ, et al. Defining the Identity and Dynamics of Adult Gastric Isthmus Stem Cells. Cell Stem Cell 2019;25:342–56.

24 Barker N, Huch M, Kujala P, et al. Lgr5(+ve) stem cells drive self-renewal in the stomach and build long-lived gastric units in vitro. Cell Stem Cell 2010;6:25–36.

25 Stange DE, Koo BK, Huch M, et al. Differentiated Troy+ chief cells act as reserve stem cells to generate all lineages of the stomach epithelium. Cell 2013;155:357–68.

26 Leushacke M, Tan SH, Wong A, et al. Lgr5-expressing chief cells drive epithelial regeneration and cancer in the oxyntic stomach. Nat Cell Biol 2017;19:774–86.

27 Sugimoto S, Ohta Y, Fujii M, et al. Reconstruction of the Human Colon Epithelium In Vivo. Cell Stem Cell 2018;22:171–6 e5.

28 Yan KS, Gevaert O, Zheng GXY, et al. Intestinal Enteroendocrine Lineage Cells Possess Homeostatic and Injury-Inducible Stem Cell Activity. Cell Stem Cell 2017;21:78–90.e6.

29 Chen B, Scurrah CR, McKinley ET, et al. Differential pre-malignant programs and microenvironment chart distinct paths to malignancy in human colorectal polyps. Cell 2021;184:6262–80.e26.

30 Qin X, Cardoso Rodriguez F, Sufi J, et al. An oncogenic phenoscape of colonic stem cell polarization. Cell 2023;186:5554–68.e18.

31 Moorman AR, Cambuli F, Benitez EK, et al. Progressive plasticity during colorectal cancer metastasis. bioRxiv 2023.

32 Karlsson M, Zhang C, Mear L, et al. A single-cell type transcriptomics map of human tissues. Sci Adv 2021;7.

33 Waddington CH. The Strategy of the Genes (1st ed.). Routledge 1957.

34 Mustata RC, Vasile G, Fernandez-Vallone V, et al. Identification of Lgr5-independent spheroid-generating progenitors of the mouse fetal intestinal epithelium. Cell Rep 2013;5:421–32.

35 Park YK, Franklin JL, Settle SH, et al. Gene expression profile analysis of mouse colon embryonic development. Genesis 2005;41:1–12.

36 Gregorieff A, Liu Y, Inanlou MR, et al. Yap-dependent reprogramming of Lgr5(+) stem cells drives intestinal regeneration and cancer. Nature 2015;526:715–8.

37 Nusse YM, Savage AK, Marangoni P, et al. Parasitic helminths induce fetal-like reversion in the intestinal stem cell niche. Nature 2018;559:109–13.

38 Yui S, Azzolin L, Maimets M, et al. YAP/TAZ-Dependent Reprogramming of Colonic Epithelium Links ECM Remodeling to Tissue Regeneration. Cell Stem Cell 2018;22:35–49 e7.

39 Kumar V, Ramnarayanan K, Sundar R, et al. Single-Cell Atlas of Lineage States, Tumor Microenvironment, and Subtype-Specific Expression Programs in Gastric Cancer. Cancer Discov 2022;12:670–91.

40 Lin R, Li C, Liu Z, et al. Genome-wide DNA methylation profiling identifies epigenetic signatures of gastric cardiac intestinal metaplasia. J Transl Med 2020;18:292.

41 Rau TT, Rogler A, Frischauf M, et al. Methylation-dependent activation of CDX1 through NF-kappaB: a link from inflammation to intestinal metaplasia in the human stomach. Am J Pathol 2012;181:487–98.

42 Kim HJ, Seo EH, Bae DH, et al. Methylation of the CDX2 promoter in Helicobacter pylori-infected gastric mucosa increases with age and its rapid demethylation in gastric tumors is associated with upregulated gene expression. Carcinogenesis 2020;41:1341–52.

43 Padmanabhan N, Kyon HK, Boot A, et al. Highly recurrent CBS epimutations in gastric cancer CpG island methylator phenotypes and inflammation. Genome Biol 2021;22:167.

44 Wang Z, Shen D, Parsons DW, et al. Mutational analysis of the tyrosine phosphatome in colorectal cancers. Science 2004;304:1164–6.

45 Lim B, Kim C, Kim JH, et al. Genetic alterations and their clinical implications in gastric cancer peritoneal carcinomatosis revealed by whole-exome sequencing of malignant ascites. Oncotarget 2016;7:8055–66.

46 Wang J, Kang WM, Yu JC, et al. Cadherin-17 induces tumorigenesis and lymphatic metastasis in gastric cancer through activation of NFkappaB signaling pathway. Cancer Biol Ther 2013;14:262–70.

47 Tong Y, Cheng PSW, Or CS, et al. Escape from cell-cell and cell-matrix adhesion dependence underscores disease progression in gastric cancer organoid models. Gut 2023;72:242–55.

48 Liao X, Wang W, Yu B, et al. Thrombospondin-2 acts as a bridge between tumor extracellular matrix and immune infiltration in pancreatic and stomach adenocarcinomas: an integrative pan-cancer analysis. Cancer Cell Int 2022;22:213.

49 Filippou PS, Karagiannis GS, Constantinidou A. Midkine (MDK) growth factor: a key player in cancer progression and a promising therapeutic target. Oncogene 2020;39:2040–54.

50 Fico F, Santamaria-Martínez A. TGFBI modulates tumour hypoxia and promotes breast cancer metastasis. Mol Oncol 2020;14:3198–210.

51 Yang S, Yang C, Yu F, et al. Endoplasmic reticulum resident oxidase ERO1-Lalpha promotes hepatocellular carcinoma metastasis and angiogenesis through the S1PR1/STAT3/VEGF-A pathway. Cell Death Dis 2018;9:1105.

52 Takei N, Yoneda A, Sakai-Sawada K, et al. Hypoxia-inducible ERO1alpha promotes cancer progression through modulation of integrin-beta1 modification and signalling in HCT116 colorectal cancer cells. Sci Rep 2017;7:9389.

53 Sun D, Guan X, Moran AE, et al. Identifying phenotype-associated subpopulations by integrating bulk and single-cell sequencing data. Nat Biotechnol 2022;40:527–38.

54 Tetteh PW, Basak O, Farin HF, et al. Replacement of Lost Lgr5-Positive Stem Cells through Plasticity of Their Enterocyte-Lineage Daughters. Cell Stem Cell 2016;18:203–13.

55 Buczacki SJ, Zecchini HI, Nicholson AM, et al. Intestinal label-retaining cells are secretory precursors expressing Lgr5. Nature 2013;495:65–9.

56 van Es JH, Sato T, van de Wetering M, et al. Dll1+ secretory progenitor cells revert to stem cells upon crypt damage. Nat Cell Biol 2012;14:1099–104.

57 Tian H, Biehs B, Warming S, et al. A reserve stem cell population in small intestine renders Lgr5-positive cells dispensable. Nature 2011;478:255–9.

58 Alvarez-Varela A, Novellasdemunt L, Barriga FM, et al. Mex3a marks drug-tolerant persister colorectal cancer cells that mediate relapse after chemotherapy. Nat Cancer 2022;3:1052–70.

59 Ganesh K, Basnet H, Kaygusuz Y, et al. L1CAM defines the regenerative origin of metastasis-initiating cells in colorectal cancer. Nat Cancer 2020;1:28–45.

60 Han T, Goswami S, Hu Y, et al. Lineage Reversion Drives WNT Independence in Intestinal Cancer. Cancer Discov 2020;10:1590–609.

61 Riera KM, Jang B, Min J, et al. Trop2 is upregulated in the transition to dysplasia in the metaplastic gastric mucosa. J Pathol 2020;251:336–47.

62 Klampatsa A, Dimou V, Albelda SM. Mesothelin-targeted CAR-T cell therapy for solid tumors. Expert Opin Biol Ther 2021;21:473–86.

63 Heinz MC, Peters NA, Oost KC, et al. Liver Colonization by Colorectal Cancer Metastases Requires YAP-Controlled Plasticity at the Micrometastatic Stage. Cancer Res 2022;82:1953–68.

## References

1 Bartfeld S, Bayram T, van de Wetering M, et al. In vitro expansion of human gastric epithelial stem cells and their responses to bacterial infection. Gastroenterology 2015;148:126–36.

2 Sato T, Stange DE, Ferrante M, et al. Long-term expansion of epithelial organoids from human colon, adenoma, adenocarcinoma, and Barrett’s epithelium. Gastroenterology 2011;141:1762–72.

3 Barker N, van Es JH, Kuipers J, et al. Identification of stem cells in small intestine and colon by marker gene Lgr5. Nature 2007;449:1003–7.

4 Yan HHN, Siu HC, Law S, et al. A Comprehensive Human Gastric Cancer Organoid Biobank Captures Tumor Subtype Heterogeneity and Enables Therapeutic Screening. Cell Stem Cell 2018;23:882–97.

5 Yan HHN, Siu HC, Ho SL, et al. Organoid cultures of early-onset colorectal cancers reveal distinct and rare genetic profiles. Gut 2020;69:2165–79.

6 Dobin A, Davis CA, Schlesinger F, et al. STAR: ultrafast universal RNA-seq aligner. Bioinformatics 2013;29:15–21.

7 Li B, Dewey CN. RSEM: accurate transcript quantification from RNA-Seq data with or without a reference genome. BMC Bioinformatics 2011;12:323.

8 de Hoon MJ, Imoto S, Nolan J, et al. Open source clustering software. Bioinformatics 2004;20:1453–4.

9 Saldanha AJ. Java Treeview--extensible visualization of microarray data. Bioinformatics 2004;20:3246–8.

10 Gremel G, Wanders A, Cedernaes J, et al. The human gastrointestinal tract-specific transcriptome and proteome as defined by RNA sequencing and antibody-based profiling. J Gastroenterol 2015;50:46–57.

11 Yu NY, Hallstrom BM, Fagerberg L, et al. Complementing tissue characterization by integrating transcriptome profiling from the Human Protein Atlas and from the FANTOM5 consortium. Nucleic Acids Res 2015;43:6787–98.

12 McCarthy DJ, Campbell KR, Lun AT, et al. Scater: pre-processing, quality control, normalization and visualization of single-cell RNA-seq data in R. Bioinformatics 2017;33:1179–86.

13 Pont F, Tosolini M, Fournie JJ. Single-Cell Signature Explorer for comprehensive visualization of single cell signatures across scRNA-seq datasets. Nucleic Acids Res 2019;47:e133.

14 Haghverdi L, Lun ATL, Morgan MD, et al. Batch effects in single-cell RNA-sequencing data are corrected by matching mutual nearest neighbors. Nat Biotechnol 2018;36:421–7.

15 Stuart T, Butler A, Hoffman P, et al. Comprehensive Integration of Single-Cell Data. Cell 2019;177:1888–902.

16 Han X, Zhou Z, Fei L, et al. Construction of a human cell landscape at single-cell level. Nature 2020;581:303–9.

17 Han S, Fink J, Jorg DJ, et al. Defining the Identity and Dynamics of Adult Gastric Isthmus Stem Cells. Cell Stem Cell 2019;25:342–56.

18 Coifman RR, Lafon S, Lee AB, et al. Geometric diffusions as a tool for harmonic analysis and structure definition of data: diffusion maps. Proc Natl Acad Sci U S A 2005;102:7426–31.

19 Angerer P, Haghverdi L, Buttner M, et al. destiny: diffusion maps for large-scale single-cell data in R. Bioinformatics 2016;32:1241–3.

20 Cao J, Spielmann M, Qiu X, et al. The single-cell transcriptional landscape of mammalian organogenesis. Nature 2019;566:496–502.

21 Bergen V, Lange M, Peidli S, et al. Generalizing RNA velocity to transient cell states through dynamical modeling. Nat Biotechnol 2020;38:1408–14.

23 Chen B, Scurrah CR, McKinley ET, et al. Differential pre-malignant programs and microenvironment chart distinct paths to malignancy in human colorectal polyps. Cell 2021;184:6262–80.e26.

24 Mustata RC, Vasile G, Fernandez-Vallone V, et al. Identification of Lgr5-independent spheroid-generating progenitors of the mouse fetal intestinal epithelium. Cell Rep 2013;5:421–32.

25 Park YK, Franklin JL, Settle SH, et al. Gene expression profile analysis of mouse colon embryonic development. Genesis 2005;41:1–12.

26 Nusse YM, Savage AK, Marangoni P, et al. Parasitic helminths induce fetal-like reversion in the intestinal stem cell niche. Nature 2018;559:109–13.

27 Yui S, Azzolin L, Maimets M, et al. YAP/TAZ-Dependent Reprogramming of Colonic Epithelium Links ECM Remodeling to Tissue Regeneration. Cell Stem Cell 2018;22:35–49 e7.

28 Gregorieff A, Liu Y, Inanlou MR, et al. Yap-dependent reprogramming of Lgr5(+) stem cells drives intestinal regeneration and cancer. Nature 2015;526:715–8.

29 Karlsson M, Zhang C, Mear L, et al. A single-cell type transcriptomics map of human tissues. Sci Adv 2021;7.

30 Ramos Zapatero M, Tong A, Opzoomer JW, et al. Trellis tree-based analysis reveals stromal regulation of patient-derived organoid drug responses. Cell 2023;186:5606–19 e24.

31 Qin X, Cardoso Rodriguez F, Sufi J, et al. An oncogenic phenoscape of colonic stem cell polarization. Cell 2023;186:5554–68.e18.

32 Volz EM, Frost SDW. Scalable relaxed clock phylogenetic dating. Virus Evolution 2017;3.

33 Kumar V, Ramnarayanan K, Sundar R, et al. Single-Cell Atlas of Lineage States, Tumor Microenvironment, and Subtype-Specific Expression Programs in Gastric Cancer. Cancer Discov 2022;12:670–91.

34 Liberzon A, Birger C, Thorvaldsdottir H, et al. The Molecular Signatures Database (MSigDB) hallmark gene set collection. Cell Syst 2015;1:417–25.

35 Sun D, Guan X, Moran AE, et al. Identifying phenotype-associated subpopulations by integrating bulk and single-cell sequencing data. Nat Biotechnol 2022;40:527–38.

36 Smyth GK. Linear models and empirical bayes methods for assessing differential expression in microarray experiments. Stat Appl Genet Mol Biol 2004;3:Article3.

37 Yang Z, Jones A, Widschwendter M, et al. An integrative pan-cancer-wide analysis of epigenetic enzymes reveals universal patterns of epigenomic deregulation in cancer. Genome Biol 2015;16:140.

38 Li H, Durbin R. Fast and accurate short read alignment with Burrows-Wheeler transform. Bioinformatics 2009;25:1754–60.

39 Li H, Handsaker B, Wysoker A, et al. The Sequence Alignment/Map format and SAMtools. Bioinformatics 2009;25:2078–9.

40 McKenna A, Hanna M, Banks E, et al. The Genome Analysis Toolkit: a MapReduce framework for analyzing next-generation DNA sequencing data. Genome Res 2010;20:1297–303.

41 DePristo MA, Banks E, Poplin R, et al. A framework for variation discovery and genotyping using next-generation DNA sequencing data. Nature genetics 2011;43:491–8.

42 Van der Auwera GA, Carneiro MO, Hartl C, et al. From FastQ data to high confidence variant calls: the Genome Analysis Toolkit best practices pipeline. Curr Protoc Bioinformatics 2013;43:11 0 1-0 33.

43 Saunders CT, Wong WS, Swamy S, et al. Strelka: accurate somatic small-variant calling from sequenced tumor-normal sample pairs. Bioinformatics 2012;28:1811–7.

44 Shen R, Seshan VE. FACETS: allele-specific copy number and clonal heterogeneity analysis tool for high-throughput DNA sequencing. Nucleic Acids Res 2016;44:e131.

